# Contextual Prediction Tunes the Tempo of Speech Segmentation

**DOI:** 10.64898/2026.03.31.713600

**Authors:** O. Platonova, O. Dogonasheva, A-L. Giraud, S. Bouton

## Abstract

Real-time speech comprehension draws on two mechanisms usually studied apart: bottom-up parsing of syllable-sized units by theta-range neural activity, and top-down prediction of next tokens based on context. Combining behavior and neurocomputational modeling, we asked how context informs syllable parsing. Human listeners (n=110) transcribed spoken sentences with altered temporal structure. Sentences were time-compressed threefold, then spread out again by interspacing silences. In a first experiment, silent gaps were either placed at natural syllable boundaries or interspaced with arbitrarily segmented speech chunks at fixed intervals. In a second experiment, gaps were placed at syllable boundaries and syllables delivered either at their natural pace (quasi-periodic) or at fixed intervals (periodic). Recognition peaked at presentation rates between 8 and 13 Hz, above the canonical *theta*-syllable range, and alignment to the syllabic structure outperformed time-based segmentation. The contextual predictability of each word, estimated by a language model, predicted recognition only when natural syllable boundaries and timing were preserved. This sensitivity to predictability was reproduced in a computational model where word-to-syllable prediction is gated by a (top-down) beta rhythm. Speech can thus be understood under heavy temporal degradation, and beyond the canonical *theta-syllable* range, if syllable timing naturalness allows context to be used.

---

Speech unfolds as a continuous stream in which sensory evidence is inherently fleeting, yet listeners achieve stable comprehension at remarkable speed. This stability implicates at least two distinct computational demands: the stream must be parsed into linguistically meaningful units (e.g., syllables and words) and contextual predictions about upcoming input must be continuously generated and integrated. These two processes, temporal segmentation and predictive inference, are frequently treated as independent, but comprehension under conditions of temporal pressure requires their active coordination. The central question is therefore not how the brain tracks speech timing, but how the speech temporal structure allows contextual prediction to act upon it.

A leading account posits that segmentation is supported by flexible theta-range dynamics (∼4-8 Hz) that align to salient acoustic landmarks (notably syllable onsets), adapting to natural variability in syllable timing rather than imposing a fixed metronome on the signal (Hovsepyan et al., 2020; Hyafil et al., 2015). On this view, acoustic evidence is accumulated over a window that opens and closes with each syllable. The theta account predicts that comprehension improves when the parse achieves this alignment. Knowing when a syllable begins and ends, however, is not knowing which syllable it is, nor how it maps onto a lexical representation. Rhythmic parsing can therefore facilitate segmentation without constituting a sufficient condition for comprehension, a limitation that becomes acute when the stream accelerates and the time available for evidence accumulation within each window is severely curtailed.

What timing leaves undermined, prior context can often supply. Predictive processing frameworks formalize comprehension as continuous probabilistic inference, integrating bottom-up sensory evidence with top-down expectations generated across representational levels (Heilbron et al., 2022; Kuperberg & Jaeger, 2016; Pickering & Gambi, 2018). On this view, temporal parsing is not only driven by external rhythm but is jointly shaped by internal generative models that influence how incoming information is segmented and interpreted over time (Arnal & Giraud, 2012; Morillon et al., 2015; ten Oever & Martin, 2021). Comprehension can then remain accurate where prior context strongly restricts the set of viable lexical candidates, even when temporal cues are degraded or the delivery rate is high. We quantify the strength of that constraint as entropy, the uncertainty a language model has about the next word given those before it: low when context is constraining, high when it is not (Hale, 2016; Linzen & Jaeger, 2016; Weissbart & Martin, 2024). A listener who exploits context should therefore recognise more of what follows where entropy is low.

Beta band (∼12–30 Hz) activity is a strong candidate for channeling top-down predictions to lower levels. In the microcircuit, feedforward prediction errors and feedback predictions occupy distinct frequency channels, the feedback component falling in the beta range (Arnal & Giraud, 2012; Bastos et al., 2015; Fontolan et al., 2014; Michalareas et al., 2016). Beta activity also reflects the maintenance and reactivation of endogenous, content-specific representations rather than stimulus-driven processing (Engel & Fries, 2010; Hovsepyan et al., 2023; Spitzer & Haegens, 2017), which is what a lexical expectation demands: it must be held before it can be brought to bear on the input. The same predictive role has been documented in language comprehension, where beta dynamics have been tied to sentence-level prediction (Lewis & Bastiaansen, 2015; Lewis et al., 2016) and to the neural tracking of linguistic structure in connected speech (Weissbart & Martin, 2024; Zioga et al., 2023). Most pertinently here, beta-band activity in time-compressed speech dissociates from theta-band syllable tracking and follows comprehension rather than acoustics (Pefkou et al., 2017).

If comprehension depends on coordinating two partially distinct inferences (namely estimating when informative units occur through temporal segmentation and what those units are likely to be through contextual prediction), then a critical test is to selectively perturb the temporal structure while holding linguistic content constant. Under conditions of high informational density, the time available for evidence accumulation within each segmentation window is reduced, potentially exposing the limits of what temporal structure alone can deliver and increasing reliance on contextual cues. Time-compressed speech provides a controlled way to challenge speech timing by increasing delivery rate while preserving the underlying linguistic sequence (Borges et al., 2018; Ghitza, 2014; Ghitza & Greenberg, 2009; Hincapié Casas et al., 2021; Pefkou et al., 2017). Intelligibility is broadly maintained under moderate compression but drops sharply at triple speed, where the time available for evidence accumulation within each segment becomes critically limited (Dupoux & Green, 1997; Pallier et al., 1998; Pefkou et al., 2017). This decline has been attributed to the failure of theta-band entrainment to track syllabic structure at elevated rates (Ghitza, 2014; Pefkou et al., 2017). Predictive-processing frameworks of comprehension suggest a complementary explanation: comprehension relies not only on what is heard but also on what the listener anticipates, and some proposals go further, holding that linguistic prediction contributes to segmenting the speech stream into analyzable units (ten Oever & Martin, 2021). If prediction both supplements the signal and helps to segment it, then at high delivery rates, where rhythmic cues are less reliable guides to where the units fall (Ghitza, 2014; Pefkou et al., 2017), comprehension should depend more on the listener’s own predictions.

The limit compression imposes can be probed by giving some of the time back: intelligibility is partially restored when silent gaps are inserted to redistribute information over time (Borges et al., 2018; Bosker & Ghitza, 2018; De La Cruz-Pavía et al., 2025; Ghitza & Greenberg, 2009; Rimmele et al., 2021). Inserting silence, however, changes three independent things: how fast the speech arrives, where the cuts fall, and how the intervals are spaced. Rate is the best characterized: comprehension improves as the delivery rate approaches the upper theta boundary, consistent with the fact that timing constrains segmentation. Where the cuts fall has received less attention, since silences are typically inserted at fixed points in the compressed stream, whether or not those points coincide with syllable onsets. Spacing is contested: Ghitza and Greenberg (2009) found intelligibility to be poorer when the inserted silences were aperiodic than when they were regular, whereas in non-speech timing aperiodic sequences support performance as well as periodic ones when their timing is predictable, even though only periodicity produces entrained modulation (Solli et al., 2025). Spacing that varies with the syllable it follows and so stays predictable from the utterance rather than departing from regularity at random has not been tested. Which of the three the comprehension gains depend on, and whether contextual prediction contributes once timing alone no longer suffices, remains open.

To separate these properties, we combined high compression with silence insertion across two experiments. Experiment 1 varies the rate at which the speech arrives and where the cuts fall, contrasting cuts placed at syllable onsets with cuts at fixed points; spacing follows placement there, since syllables differ in length while fixed chunks do not. Experiment 2 removes that confound by placing every cut at a syllable onset and varying the spacing alone, comparing intervals held constant with intervals proportional to the syllable they follow, again across a range of rates. Between them the two experiments separate the three properties. Both also let us ask whether contextual predictability contributes and where. It should contribute only where segmentation delivers syllabic units, since that is the level over which lexical prediction operates: at matched delivery rate, entropy should track recognition under syllable-aligned segmentation and not under time-based segmentation. Experiment 2 sharpens the test, since units are delivered in both of its conditions: if what matters is that the timing follows those units rather than coinciding with them, entropy should track recognition under quasi-periodic pacing and not under periodic pacing.

The experiments can address whether syllabicity drives contextual predictability, but not how. We therefore implemented a computational model in which the best candidate gating mechanism, a modulated beta-band rhythm, controls the flow of word-level predictions onto syllabic processing, and compared it with the same model stripped of that gating. If the account is right, only the model with the gate in place should show the listeners’ dependence on predictability, and within that model the predictions released onto a unit should determine whether it is recognized, more so where the boundaries fall on syllables.

## RESULTS

### Bottom-up: Timing Helps Only When It Follows the Syllabic Units

#### Experiment 1. Temporal Structure and Linguistic Alignment Shape Speech Comprehension under High Compression

Experiment 1 examines how speech comprehension is affected when the temporal structure of the acoustic signal is severely distorted. All sentences are time-compressed by a factor of three, resulting in a syllabic rate of 16.1 Hz. Silent gaps are then inserted between successive chunks to reduce the effective delivery rate without restoring fine-grained acoustic detail. Varying pause duration produces **six delivery rates** (4.6, 5.4, 6.5, 8.1, 10.8, and 12.9 Hz).

Segmentation structure is independently manipulated to dissociate externally driven temporal sampling from endogenous linguistic units. In the **time-based segmentation** condition, speech is divided into uniform 62-ms chunks corresponding to the average compressed syllable duration, imposing a strictly regular temporal structure that disregards syllabic boundaries. In the **syllable-aligned segmentation** condition, chunk boundaries are placed aligned with syllable onsets, preserving access to linguistically meaningful units while retaining natural temporal variability (Figure 1A). This design allows the contribution of delivery rate and boundary alignment to be assessed independently and their interaction to be characterized under conditions of high temporal distortion.

Fifty adult participants (26 females; *mean* age = 39.3, *SD* = 9.4) each listened to 102 unique sentences drawn from the TIMIT corpus (Garofolo et al., 1993), matched for duration, syllable count, and prosodic balance. Each sentence is presented once, and participants transcribe what they understand via keyboard. Responses are aligned to the reference sentence and scored automatically, token by token, and the trial score is the proportion of tokens recognized. A token is an entry in the sub-word vocabulary of GPT-2, the language model used to compute contextual entropy: a whole word when the word is frequent, a fragment of one when it is rare.

**Figure 1.**
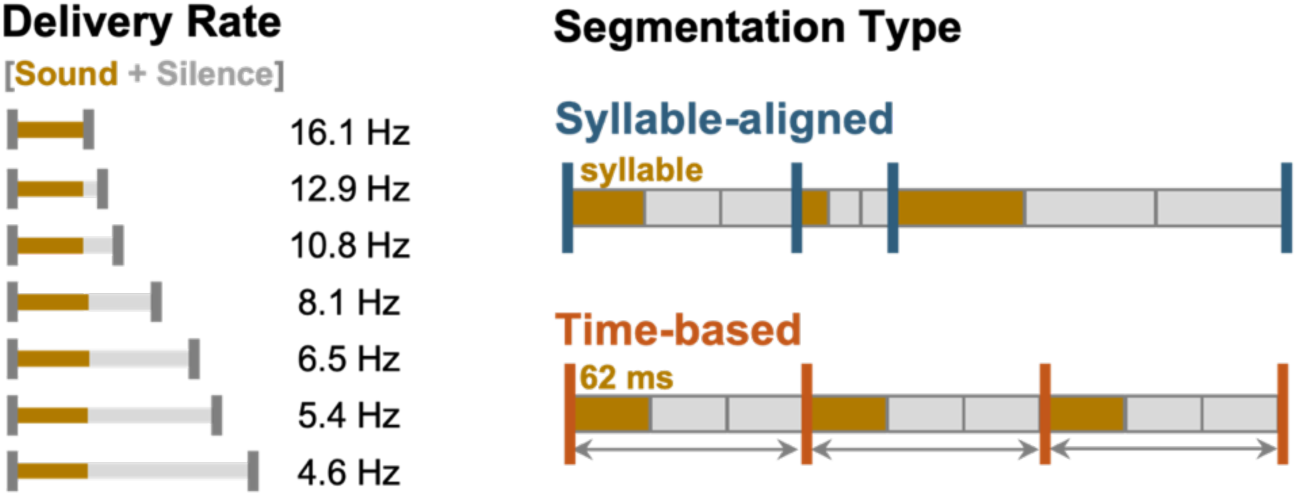
Paradigm used in Experiment 1. Sentences are compressed by a factor of 3, increasing the syllabic rate to 16.1 Hz. Silent gaps are inserted between chunks as a proportion of a sound chunk duration (adapted to each syllable) to generate six delivery rates ranging from 4.6 to 12.9 Hz. Two segmentation types are tested: syllable-aligned segmentation (top), in which speech chunks are aligned with syllable boundaries; and time-based segmentation (bottom), in which uniform 62-ms chunks are separated by pauses, irrespective of linguistic structure.

Recognition is modelled with a beta mixed-effects regression on Segmentation type x Delivery rate, with crossed random intercepts for participant and sentence. The main effect of segmentation (Figure 2C), the effect of delivery rate, and their interaction (Figure 2D) are read from this model. Delivery rate is coded as a categorical factor for comparisons at rates and as a mean-centered second-order polynomial for the shape of the rate function (Figure 2B). Type III tests use sum-to-zero contrasts so that main effects are averaged over the levels of the other factor.

Two comparisons fall outside this design: uncompressed against fully compressed speech (Figure 2A) and each segmentation type against the fully compressed baseline (Figure 2C). Neither condition carries a segmentation type, so neither can enter the factorial model, and both are estimated separately. In the analysis of the shape of the rate function (Figure 2B), the fully compressed condition is entered at its syllabic rate of 16.1 Hz, which extends the range over which the curve is estimated.

##### Severe temporal compression degrades speech comprehension

Three-fold compression produces a large decrease in recognition relative to uncompressed speech (mean difference = 51.98, 95% CI [46.25, 57.72]; *χ*^2^(1) = 77.31, *p* < .001; Figure 2A), replicating prior findings (Dupoux & Green, 1997; Pallier et al., 1998; Pefkou et al., 2017). At the fully compressed rate (16.1 Hz), listeners recover 19.3% of the materials, against 78.5% for uncompressed sentences, indicating that the compressed acoustic stream alone provides insufficient temporal support for reliable word recognition. This condition serves as the baseline against which the effects of delivery rate and segmentation structure are assessed.

##### Comprehension peaks above the canonical theta range

Averaged across segmentation conditions, recognition varies systematically with delivery rate. The primary model, refitted with delivery rate as a mean-centered second-order polynomial and with the fully compressed condition entered at its syllabic rate of 16.1 Hz, reveals a non-linear relationship (Wald *χ*^2^(2) = 117.81, *p* < .001; the quadratic term against a linear-only model, *χ*^2^(1) = 92.70, p < .001, Figure 2B). Recognition improves as delivery rate is reduced from 16.1 Hz and is highest between 8.1 and 12.9 Hz, where it is essentially flat (28.1%, 27.4%, and 27.9%), with the fitted maximum near 10.6 Hz. It then declines at slower rates, reaching 17.0% at 4.6 Hz, no better than the 19.3% obtained under the fully compressed baseline. Recognition is therefore best at rates above the canonical theta range, and slowing delivery beyond that point returns the listener to the baseline. Whether this reflects delivery rate on its own, or where the segmentation boundaries fall, is what the next analyses address.

##### Syllable-aligned segmentation consistently outperforms time-based segmentation

Averaged over delivery rates, the primary model reveals a significant main effect of segmentation type: syllable-aligned segmentation exceeds time-based segmentation by an odds ratio of 1.33 (95% CI [1.24, 1.43], z = 7.89, *χ*^2^(1) = 62.21 p < .001; Figure 2C). Set against the fully compressed baseline, that difference separates a condition that improves unsegmented compressed speech from one that does not. Syllable-aligned segmentation raises recognition substantially above the baseline (odds ratio = 1.45, 95% CI [1.23, 1.70], z = 5.40, p < .001), whereas time-based segmentation does not differ reliably from it (odds ratio = 1.10, 95% CI [0.93, 1.29], z = 1.33, p = .38). Inserting silence is therefore not sufficient on its own: recognition improves only when the cuts fall at syllable onsets. Regular pacing alone does not restore comprehension, consistent with the view that temporal cues are exploited when they map onto internal phonological representations.

**Figure 2.**
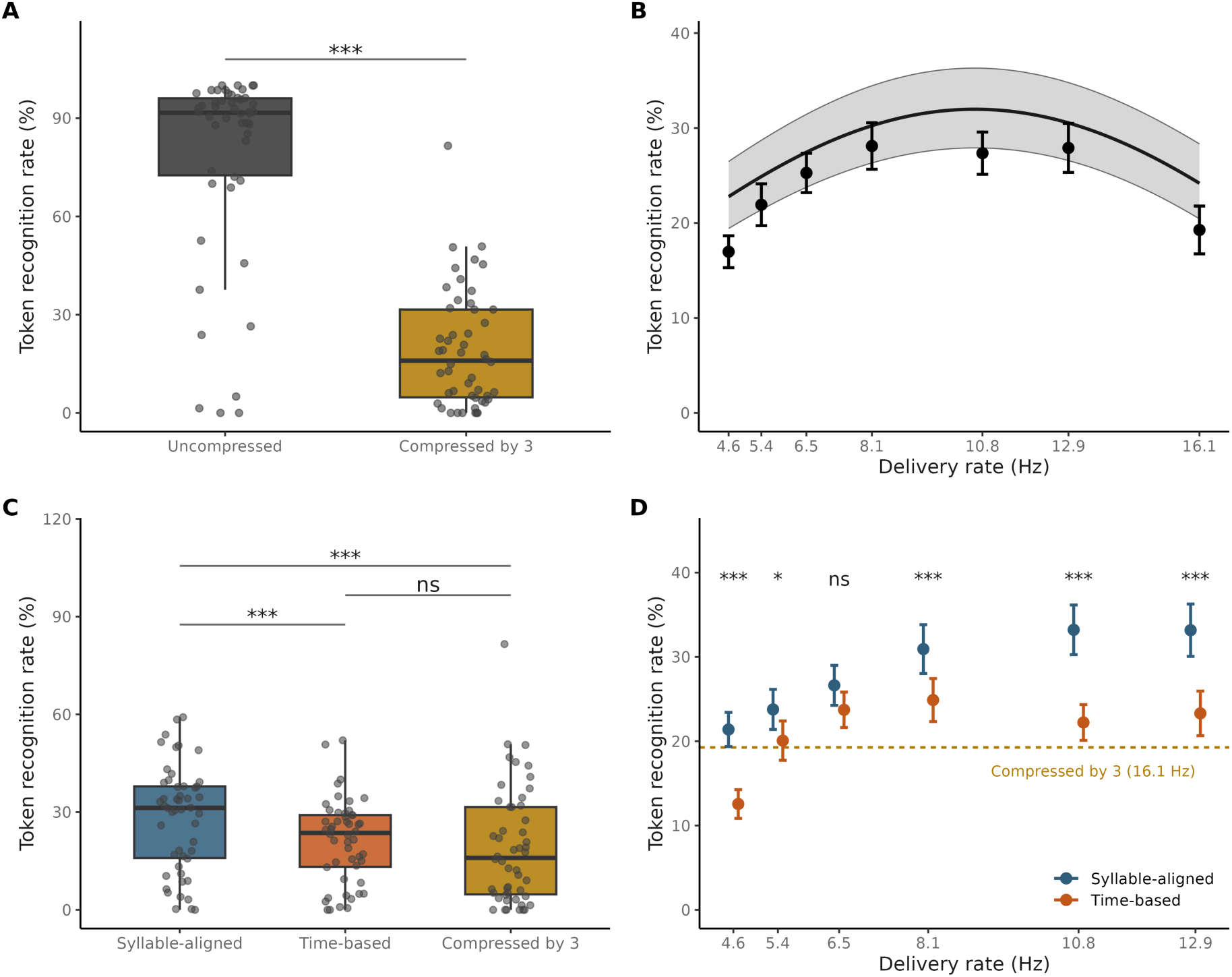
Effects of Segmentation Type and Delivery Rate on Word Recognition. Recognition is scored token by token, a token being a whole word for frequent words and a fragment of one for rarer words. (A) Effect of time compression. Three-fold compression significantly reduces the token recognition rate relative to uncompressed speech (***, p < .001). Each dot represents one participant’s mean recognition rate; boxplots indicate the median (horizontal line) and interquartile range (IQR), with whiskers extending to the most extreme values within 1.5 × IQR (Tukey convention). (B) Token recognition rate as a function of delivery rate (Hz), averaged across segmentation conditions, with the fully compressed condition plotted at its syllabic rate of 16.1 Hz. Points show the observed mean across participants (± SEM). The solid curve is the fitted quadratic beta mixed-effects model, integrated over the participant and sentence random intercepts so that it estimates the population average, and the shaded band its 95% confidence interval. (C) Token recognition rate by segmentation type. Dots and boxplots as in (A). Syllable-aligned segmentation exceeds both time-based segmentation and fully compressed speech, whereas time-based segmentation does not differ reliably from fully compressed speech (Tukey-adjusted; ns, not significant; ***, p < .001). (D) Segmentation type x delivery rate. Mean token recognition rate (± SEM) for syllable-aligned (blue) and time-based segmentation (orange). The dashed golden line marks the fully compressed baseline at 16.1 Hz. Labels give the syllable-aligned versus time-based contrast at each rate, corrected across the six rates (ns, not significant; *, p < .05; **, p < .01; ***, p < .001).

##### Syllabic alignment becomes critical outside the optimal temporal regime

To assess whether the benefit of syllable-aligned segmentation depends on delivery rate, we examine the Segmentation × Delivery Rate interaction from the primary model. The interaction does not reach significance (*χ*^2^(5) = 9.26, p = .099), so the contrasts below describe where the advantage lies (Figure 2D). Syllable-aligned segmentation outperforms time-based segmentation at faster delivery rates (8.1 Hz: odds ratio = 1.38, 95% CI [1.15, 1.64], z = 3.57, p <.001; 10.8 Hz: odds ratio = 1.49, 95% CI [1.25, 1.78], z = 4.45, p < .001; 12.9 Hz: odds ratio = 1.48, 95% CI [1.25, 1.63], z = 3.48, p < .001) and at the slowest rates (4.6 Hz: odds ratio = 1.37, 95% CI [1.15, 1.76], z = 4.48, p < .001; 5.4 Hz: odds ratio = 1.25, 95% CI [1.05, 1.48], z = 2.46, p = .017), the latter reflecting a pronounced decline in performance under time-based segmentation. At intermediate delivery rates, by contrast, segmentation effects are attenuated: no reliable difference between segmentation types is observed at 6.5 Hz (odds ratio = 1.08, 95% CI [0.91, 1.29], z = 0.88, p = .38). All contrasts are corrected across the six rates. Thus, the benefit of syllable-aligned over time-based segmentation is weakest near the center of the theta range, where stimulus-driven temporal structure appears sufficient to support comprehension regardless of boundary position. Outside this regime, whether at faster or slower rates, syllable-aligned segmentation becomes increasingly critical, revealing the limits of what timing alone can deliver.

Experiment 1 establishes that comprehension under severe temporal distortion depends jointly on delivery rate and on where segmentation boundaries fall. Recognition is best across a broad band above the canonical theta range, from 8 to 13 Hz, and declines as delivery rate slows further: at 4.6 Hz, it is no better than for fully compressed speech. Slowing the delivery rate is therefore not sufficient by itself, and beyond a certain point it stops helping altogether. Recognition improves only where segmentation boundaries coincide with syllabic units, and the advantage of syllable-aligned over time-based segmentation is smallest in the middle of the theta range and grows on either side of it.

A key limitation of Experiment 1 is that its two conditions differ on two counts at once: where the silent gaps fall and how the syllables are spaced. In the time-based condition, gaps are interspaced with arbitrarily segmented speech chunks, so they fall without regard to syllable boundaries, and the chunks arrive at fixed intervals. In the syllable-aligned condition, gaps are laced at natural syllable boundaries, and each gap is scaled to the duration of the syllable it follows, so syllables arrive at their natural pace. The advantage of syllable-aligned segmentation may therefore come from where the gaps fall, from the pace they impose, or from both.

Experiment 2 is designed to resolve this ambiguity. Silent gaps are placed at natural syllable boundaries in every condition, so their placement is held constant and only the pace varies: syllables are delivered either at their natural pace, each gap being scaled to the syllable it follows (quasi-periodic), or at fixed intervals (periodic), at four matched delivery rates. If fixed pacing is what carries the benefit, the periodic condition should do at least as well as the quasi-periodic one. If instead the benefit comes from syllables arriving at their natural pace, the quasi-periodic condition should do better.

#### Experiment 2. Temporal Regularity Modulates Comprehension under Syllable-Based Segmentation

All sentences are time-compressed by a factor of three, increasing the syllabic rate to 16.1 Hz. Silent gaps are inserted between syllables to reduce the delivery rate, producing four delivery rates (5.3, 6.4, 7.9, and 10.6 Hz). Temporal regularity is manipulated by varying how these gaps are spaced. In the *quasi-periodic* condition, each gap is proportional to the compressed syllable it follows, so syllables arrive at their natural pace, with the variability of their durations preserved. In the *periodic* condition, each gap is set to that syllables arrive at fixed intervals (Figure 3). Sixty adult participants (31 females, mean age = 44.6, *SD* = 11.6) complete the task, which is otherwise identical to Experiment 1.

**Figure 3.**
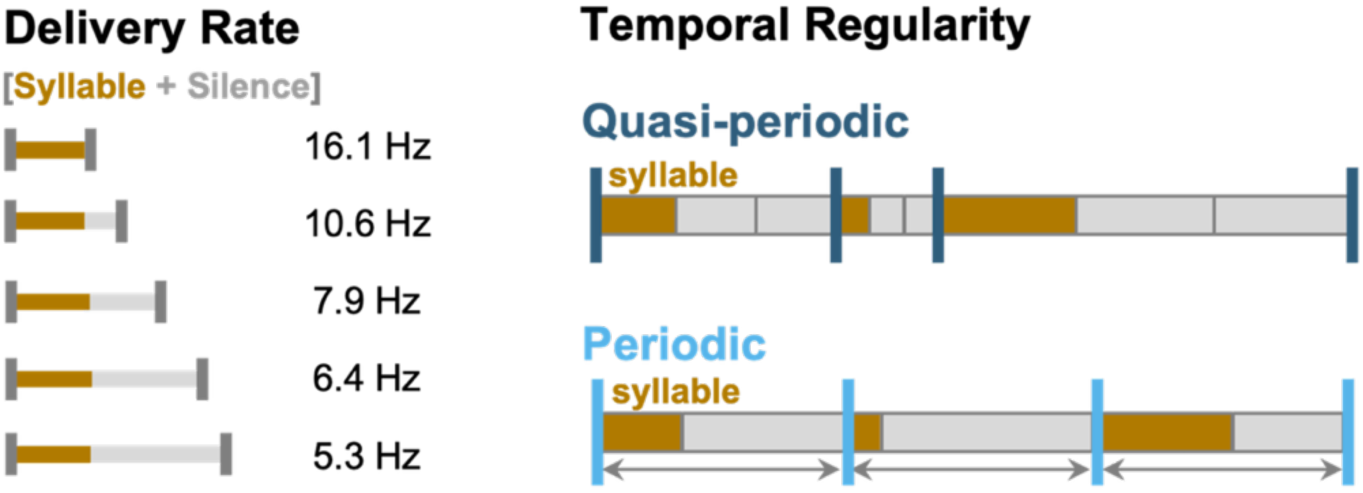
Paradigm used in Experiment 2. Sentences are compressed by a factor of three, raising the syllabic rate to 16.1 Hz, then cut at syllable boundaries. Silent gaps inserted between syllables reduce the delivery rate, giving four rates (5.3, 6.4, 7.9 and 10.6 Hz). Cuts fall at syllable onsets in each condition, so their placement is held constant and only the pace varies. In the quasi-periodic condition, the gap following each syllable lasts that syllable’s own duration multiplied by a fixed factor, 0.5, 1, 1.5 or 2, which is what sets the delivery rate. Because syllables differ in duration, the gaps vary within a sentence even though the factor does not. The example illustrated uses a factor of two. In the periodic condition, each gap is set so that syllables arrive at fixed intervals, the duration of one syllable together with the gap that follows it being matched to the median of the corresponding quasi-periodic condition, so that mean delivery rate is equated across the two conditions.

Recognition is modelled with a beta mixed-effects regression on Temporal regularity x Delivery rate, with crossed random intercepts for participant and sentence. Delivery rate is entered as a categorical factor for comparisons at particular rates, and as a mean-centered second-order polynomial for the shape of the rate function. Type III tests use sum-to-zero contrasts, as in Experiment 1. The uncompressed versus fully compressed comparison (Figure 4A) is estimated separately, since neither condition carries a temporal-regularity level and neither can enter the factorial model.

##### Inserting silent gaps between syllables improves recognition

Inserting silent gaps between syllables produces a significant increase in token recognition rate relative to the fully compressed baseline (mean difference = 57.72, 95% CI [47.97, 67.46]; *χ*^2^(1) = 49.80, *p* < .001, Figure 4A). Both periodic and quasi-periodic segmentation significantly improve comprehension relative to the unsegmented compressed condition (Periodic vs. Compressed: odds ratio = 1.53, 95% CI [1.35, 1.74], z = 7.86, p < .001; Quasi-periodic vs. Compressed: odds ratio = 1.83, 95% CI [1.61, 2.08], z = 11.16, p < .001, Figure 4B). Averaged across delivery rates, quasi-periodic timing outperforms strictly periodic timing (odds ratio = 1.20, 95% CI [1.11, 1.30], z = 5.30, p < .001). Because every cut falls at a syllable onset in both conditions, this difference is attributable to the spacing of the intervals alone: letting the intervals follow the syllables they separate helps beyond what syllable alignment provides.

##### Comprehension peaks at delivery rates above the canonical theta range

Averaged across the two regularity conditions, recognition is strongly modulated by delivery rate (Figure 4C). The primary model shows significant linear (z = 4.04, p < .001) and quadratic (z = -10.42, p < .001) effects, and the quadratic term is justified against a linear-only model (*χ*²(1) = 108.48, *p* < .001). Recognition improves as delivery slows from 16.1 Hz, is highest between 7.9 and 10.6 Hz, and declines at slower rates, so that as in Experiment 1 the optimum lies above the canonical theta range. Unlike Experiment 1, however, the shape of this profile differs between the two conditions (*χ*^2^(2) = 10.48, *p* = .005), which the next analysis takes up.

##### Temporal flexibility supports comprehension at faster delivery rates

The Temporal Regularity x Delivery Rate interaction is significant (*χ*^2^(3) = 10.51, *p* = .015, Figure 4D). Periodic and quasi-periodic timing do not differ at the slowest rates (5.3 Hz: odds ratio = 1.12, 95% CI [0.98, 1.28], z = 1.65, p = .12; 6.4 Hz: 1.11, 95% CI [0.97, 1.27], z = 1.55, p = .12). A modest advantage for quasi-periodic timing appears at 7.9 Hz (odds ratio = 1.17, 95% CI [1.02, 1.34], z = 2.31, p = .041) and becomes pronounced at 10.6 Hz (odds ratio = 1.45, 95% CI [1.27, 1.66], z = 5.51, p < .001), the four contrasts being corrected across the family. The two conditions diverge because they behave differently at the fast end: under quasi-periodic timing recognition keeps improving up to 10.6 Hz, its highest value, whereas under strictly periodic timing it turns down after 7.9 Hz. Once every boundary falls at a syllable onset, what governs recognition at fast delivery rates is therefore not how regular the timing is, but whether it follows the syllables.

Experiment 2 shows that regularity is not itself what helps. When every gap falls at a syllable boundary, syllables delivered at their natural pace are recognized better than syllables delivered at fixed intervals, and the difference widens as delivery speeds up: at 10.6 Hz, recognition keeps improving when syllables arrive at their natural pace while it falls when they arrive at fixed intervals. Taken with Experiment 1, this isolates two properties of the temporal organization that supports recognition: gaps that fall at syllable boundaries, and intervals that follow the durations of the syllables they carry.

**Figure 4.**
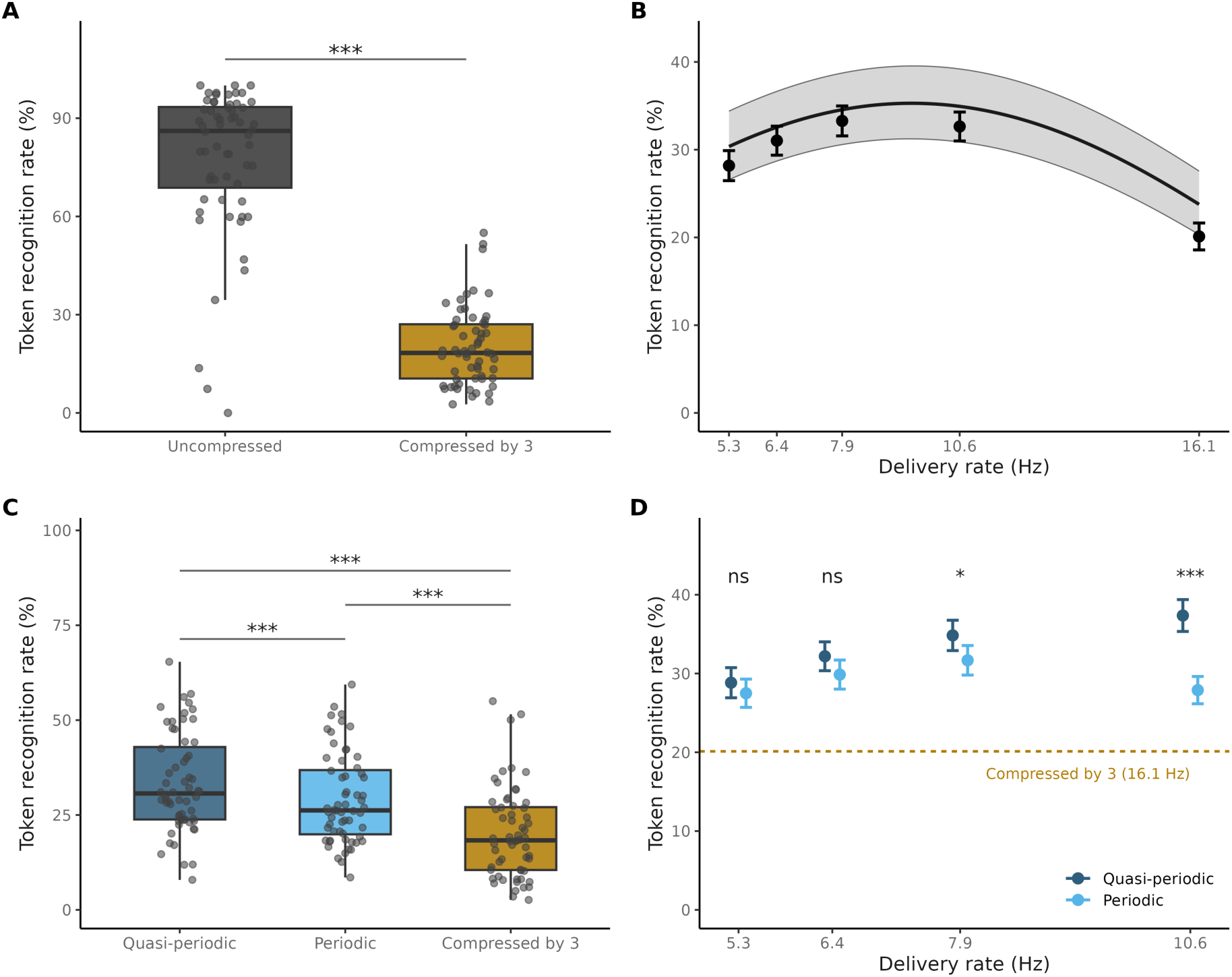
Temporal regularity and delivery rate shape token recognition. (A) Effect of time compression. Fully compressed speech (“Compressed by 3”) produces markedly lower token recognition rates than uncompressed speech (*** p < .001). Each dot represents one participant; boxplots indicate the median and the interquartile range (IQR), with whiskers extending to the most extreme values within 1.5 x IQR. (B) Effect of delivery rate. Mean recognition rate (± SEM) averaged over the two regularity conditions, with the fully compressed condition plotted at its syllabic rate of 16.1 Hz. The curve is the fitted quadratic model (± 95% CI). Recognition is highest between 7.9 and 10.6 Hz and falls at slower rates. This profile is not shared by the two conditions, whose rate functions differ in shape (see D). (C) Effect of temporal regularity. Quasi-periodic timing, in which syllables arrive at their natural pace (dark blue), periodic timing, in which they arrive at fixed intervals (light blue), and the fully compressed baseline (gold). Both regularity conditions exceed the compressed baseline, and quasi-periodic timing outperforms periodic timing overall. (D) Temporal regularity x delivery rate. Mean token recognition rate (± SEM) for periodic and quasi-periodic timing at each delivery rate. The dashed golden line marks the fully compressed baseline. Labels give the quasi-periodic versus periodic contrast at each rate, corrected across the four rates (ns, not significant; *, p < .05; ***, p < .001).

So far, this benefit has been read as a matter of parsing: place the cuts well and more words are recognized. The same arrangement may also be what allows context to act on the signal. The two accounts differ in what they predict about contextual prediction: on the first, context should help wherever recognition is possible at all; on the second, only where the timing follows the syllables. The next analysis separates them, using token-by-token predictability, quantified as the entropy of the next token given the preceding context: the higher the entropy, the less predictable what follows.

### Top-Down: Context Helps Only Where the Timing Follows the Syllables

Entropy is computed at each token position from the intact sentence, so the same value applies to every listener at a given position. It is highest at sentence onset and falls as context accumulates (Figure S1A).

Both experiments are analyzed with the same specification. Entropy and recovery are both defined at the token level, so the analysis is carried out at that level: each token is scored as correctly recognized or not and modelled with a binomial mixed-effects regression on entropy x condition x delivery rate, with crossed random intercepts for participant, sentence, trial, and token position. The first three capture the levels at which trials are sampled; the fourth absorbs the fact that tokens late in a sentence are recognized less often than early ones. What the design asks of this model is where entropy affects recognition, so the slope of recognition on entropy is estimated in each cell, and the twelve cells of Experiment 1 and the eight of Experiment 2 are corrected as single families with Benjamini-Hochberg false discovery rate.

#### Entropy predicts recognition where segmentation delivers syllabic units

In Experiment 1, the token-level analysis reproduces the bottom-up pattern of Figure 2 (segmentation: *χ*^2^(1) = 16.99, p < .001; delivery rate: *χ*^2^(5) = 37.37, p < .001; segmentation x delivery rate: *χ*^2^(5) = 21.39, p < .001). Turning to the terms of interest, higher entropy lowers the probability that a token is recognized (*χ*^2^(1) = 11.17, p < .001), so contextual predictability carries a measurable benefit even under three-fold compression. That benefit is not uniform. It varies with delivery rate (*χ*²(5) = 17.13, p = .004) and jointly with segmentation and delivery rate (*χ*²(5) = 13.27, p = .021; Figure 5). Entropy slopes are reliably negative under syllable-aligned segmentation at 4.6 Hz (slope = -0.108, z = -3.34, p = .005), 8.1 Hz (-0.109, z = -3.97, p < .001) and 10.8 Hz (-0.073, z = -2.51, p = .036), while at 12.9 Hz the slope is indistinguishable from zero (-0.012, z = -0.42, p = .67). Under time-based segmentation only 6.5 Hz reaches significance (-0.081, z = -2.68, p = 0.030), the remaining five slopes being small and unreliable.

**Figure 5.**
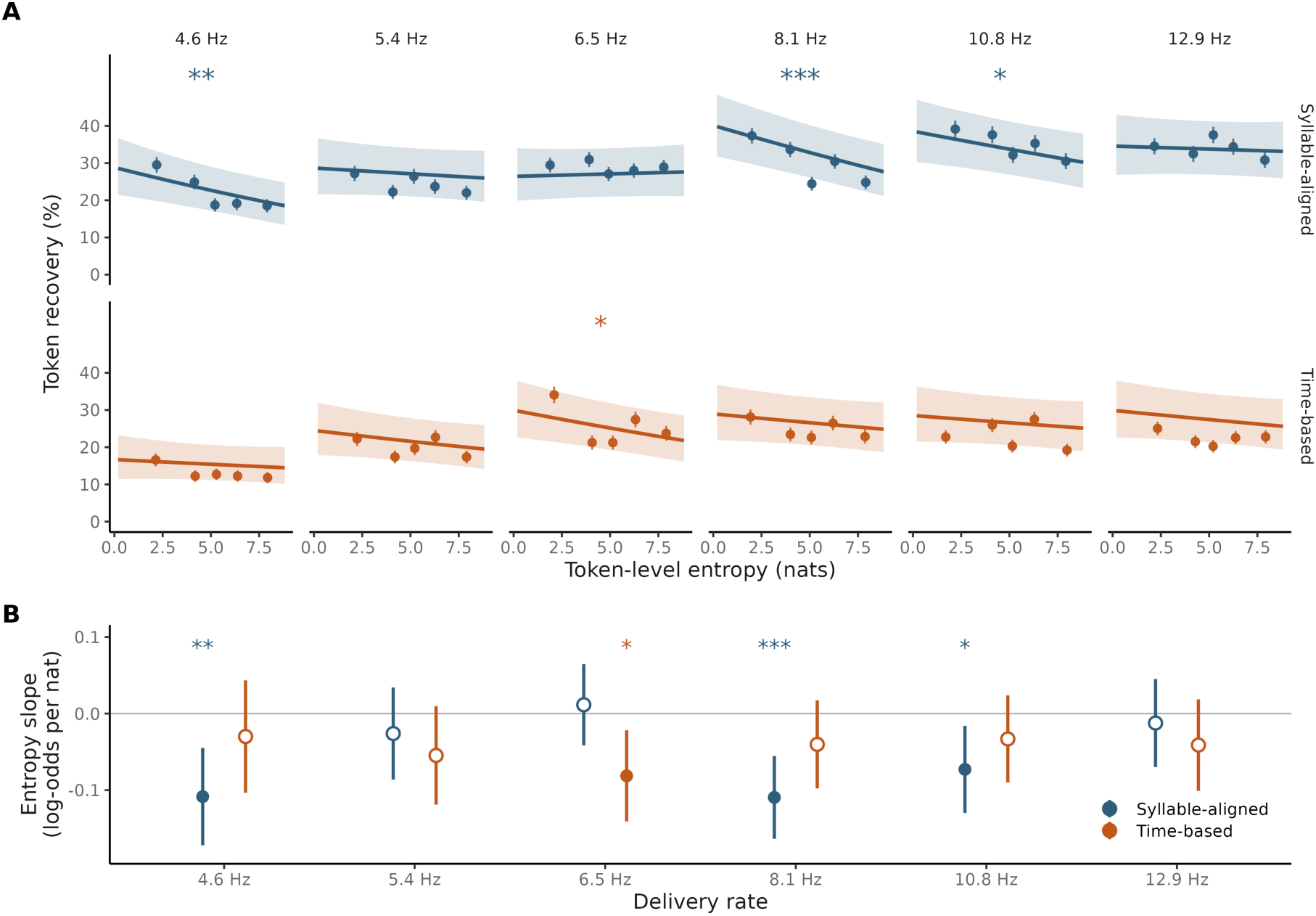
Higher entropy lowers token recovery mainly under syllable-aligned segmentation (Experiment 1). (A) Probability that a token is recovered as a function of token-level entropy, for syllable-aligned (blue, top row) and time-based segmentation (orange, bottom row), at each delivery rate (columns). Points show observed proportions within five entropy bins per cell (± SEM); the line shows the fitted binomial mixed-effects model, and the band its 95% confidence interval. (B) Entropy slopes (log-odds of token recovery) with 95 % confidence intervals for each segmentation type at each delivery rate. Filled points mark slopes that remain significant after Benjamini-Hochberg correction across the twelve cells, open points mark those that do not (*, p < .05; **, p < .01; ***, p < .001).

#### Entropy predicts recognition under quasi-periodic pacing

In Experiment 2, the token-level analysis reproduces the bottom-up pattern of Figure 4 (delivery rate: *χ*^2^(3) = 26.50, p < .001; regularity: *χ*^2^(1) = 2.92, p = .087; regularity x delivery rate: *χ*^2^(3) = 17.09, p < .001). As in Experiment 1, higher entropy lowers the probability that a token is recovered (*χ*^2^(1) = 5.93, p = .015), and the dependence on how the signal is laid out in time does not reach significance here (entropy x regularity x delivery rate, *χ*^2^(3) = 7.37, p = .061; Figure 6). Under quasi-periodic pacing, where each silent gap is proportional to the syllable it follows, entropy predicts recognition at every rate tested, with p values corrected across the eight cells: 5.3 Hz (slope = -0.049, z = - 2.43, p = .030), 6.4 Hz (-0.053, z = -2.62, p = .024), 7.9 Hz (-0.099, z = -5.03, p < .001) and 10.6 Hz (-0.068, z = -3.55, p = .002). Under strictly periodic pacing, no slope reaches significance at any rate (all p > .19).

**Figure 6.**
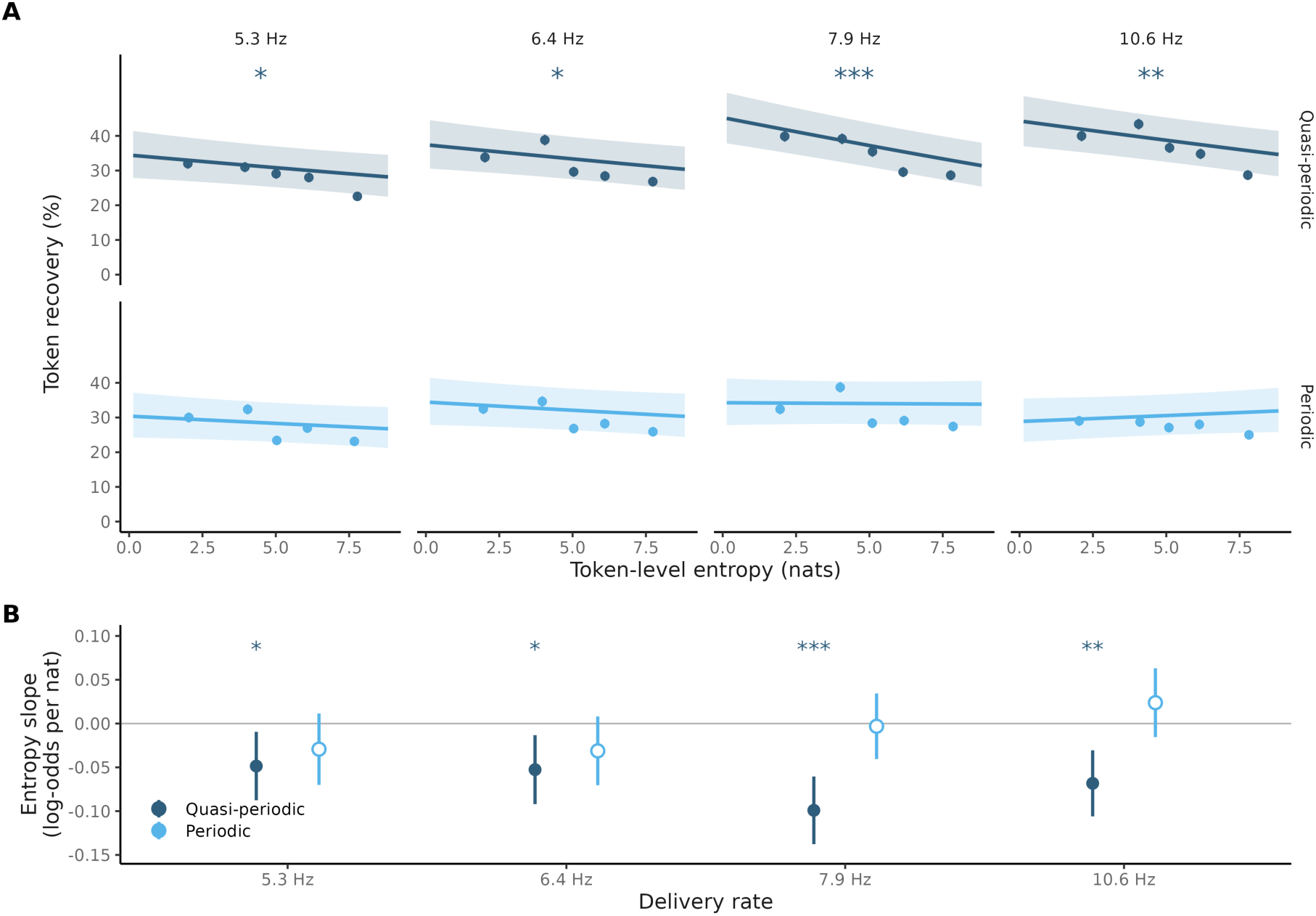
Higher entropy lowers token recognition under quasi-periodic timing at every delivery rate (Experiment 2). (A) Probability that a token is recognized as a function of token-level entropy, for quasi-periodic (dark blue, top row) and periodic segmentation (light blue, bottom row), at each delivery rate (column). Points show observed proportions within five entropy bins per cell (± SEM); the line shows the fitted binomial mixed-effects model, marginalized over the random effects, and the band its 95% confidence interval. Asterisks mark cells whose entropy slope differs from zero after correction. (B) Entropy slopes (change in the log-odds of token recovery per unit of entropy) with 95% confidence intervals. Filled points mark slopes that remain significant after Benjamini-Hochberg correction across the eight cells, open points mark those that do not (*, p < .05; **, p < .01; ***, p < .001).

Taken together, the two experiments point to a single condition under which contextual predictability bears on recognition: the stream must deliver syllables, and deliver them at intervals that follow the durations of those syllables rather than a constant period. Every condition tested falls on one side or the other of that criterion, and the entropy effect follows it. Syllable-aligned segmentation in Experiment 1 and quasi-periodic pacing in Experiment 2 satisfy both terms, and the effect is present in both, with slopes of the same magnitude at the rates the two experiments share. Strictly periodic pacing satisfies the first and not the second, since every cut falls at a syllable onset while the interval between cuts is constant, and there the effect is absent at every rate. Time-based segmentation satisfies neither, except at 6.5 Hz, where each 62-ms chunk carries one compressed syllable and the chunks arrive at close to a natural syllable rate: that is where the effect appears on that side, and the next largest slope is at 5.4 Hz, the other rate whose chunks approach a syllable in duration.

What the experiments do not show is how a prediction is held until the moment at which it can be applied. That calls for a mechanism in which the content of a prediction and its timing are handled together, and beta-band dynamics are the candidate. Indeed, beta carries the feedback component of cortical prediction, and it tracks the maintenance of content-specific representations rather than the stimulus itself. We therefore implemented a model in which a beta-band gate releases token-level predictions onto syllabic parsing, and compared it with the same model with the gate help open so that predictions are released continuously and at full strength. Applied to the conditions of Experiment 1, the comparison asks whether that gate is what makes recognition follow contextual predictability, and whether it does so only where the boundaries fall on syllables.

#### Computational Modeling

The goal of the computational modeling is to assess whether the sensitivity patterns observed in the listeners can arise from a principled predictive account rather than from rhythmic alignment alone. The behavioral influence of contextual predictability, indexed by token entropy, is formalized as β-mediated lexical prediction: word-level representations generate top-down expectations over syllabic transitions, and the beta rhythm gates the degree to which these expectations shape syllabic parsing. The manipulations of Experiments 1 and 2 are applied to the β-BRyBI model (Dogonasheva et al., 2025), a hierarchical generative architecture in which higher-level lexical states generate predictions over lower-level acoustic representations through precision-weighted message passing (Figure S3). Contextual prediction is here an internally generated expectation, not an externally driven rhythmic process.

In β-BRyBI, syllables are the representational units that organize the hierarchy. Incoming speech is parsed through a flexible theta-paced sampling process that adapts to follow the syllabic rhythm of the acoustic signal, supplying candidate syllabic units to the lexical level, and lexical states send prediction back over syllabic identity. The model has no token level: the token enters only at measurement stage, where model output, listener responses and contextual entropy are read on the same byte-pair units. Two consequences follow from this architecture, and the experimental contrasts were designed to test them. If predictions need syllabic units to apply to, their influence should depend on whether segmentation delivers such units, and should weaken when boundaries are artificially placed on a clock. And if predictions specify which syllabic unit is likely to come next rather than when it will arrive, their influence should not depend on the regularity of the intervals between boundaries.

The model runs in two states. In β-ON, the top-down gain is modulated in the beta range at 17 Hz, so that the strength with which a prediction is released varies from moment to moment. In β-OFF that top-down gain is held constant at its maximum, so predictions remain available throughout: β-OFF removes the beta modulation, not the top-down pathway. All model output reads at the level of the token, the byte-pair unit over which the language model defines its vocabulary, so that model and listeners are scored on the same units. The two sides are compared on the same recordings and the same tokens, the first token of each sentence being excluded on both for want of preceding context. Recognition is modelled with a binomial family and a logit link, with random intercepts for sentence, token position, and run, and omnibus effects are tested by Type III Wald chi-square under sum-to-zero contrasts.

#### **β**-BRyBI reproduces the two experimental effects and matches listeners within the theta range

Segmentation type and delivery rate in Experiment 1, temporal regularity and delivery rate in Experiment 2, act on the model as they act on the listeners. In Experiment 1 the model recognizes more tokens under syllable-aligned than under time-based segmentation (*χ*^2^(1) = 127.24, OR = 2.24, 95% CI [1.94, 2.57], z = 11.28), and recognition varies with the delivery rate (*χ*^2^(5) = 135.98). In Experiment 2 it recognizes more tokens under quasi-periodic than under periodic timing (*χ*^2^(1) = 11.21, OR = 1.37 [1.14, 1.64], z = 3.35), with an effect of rate (*χ*^2^(3) = 141.39) and an interaction between the two (*χ*^2^(3) = 84.48), again as in the listeners.

Model and listeners are then compared directly, the two states pooled. The distance between them depends on delivery rate (*χ*^2^(5) = 144.69 in Experiment 1 and *χ*^2^(3) = 90.80 in Experiment 2). Within the theta range the model reaches the listeners’ level in all four conditions, recognizing tokens at 0.83 times their odds under time-based segmentation, 0.92 under quasi-periodic timing and 0.76 under periodic timing, and at 1.43 under syllable-aligned segmentation. Above 8 Hz, it falls short in all four conditions, at 0.30 to 0.47 times their odds (all p ≤ 1.3 × 10^−4^) and in every condition the shortfall is larger above that limit than below it (all p ≤ 0.0022) (Figure 7A). Thus, within the theta range, reaching human level across four conditions that differ from one another by fifteen to twenty points of recognition is a strong constraint, and it places the model on the structure the listeners use. Outside that range, the model no longer keeps up, so its account of segmentation is bounded by the rate at which it parses.

**Figure 7.**
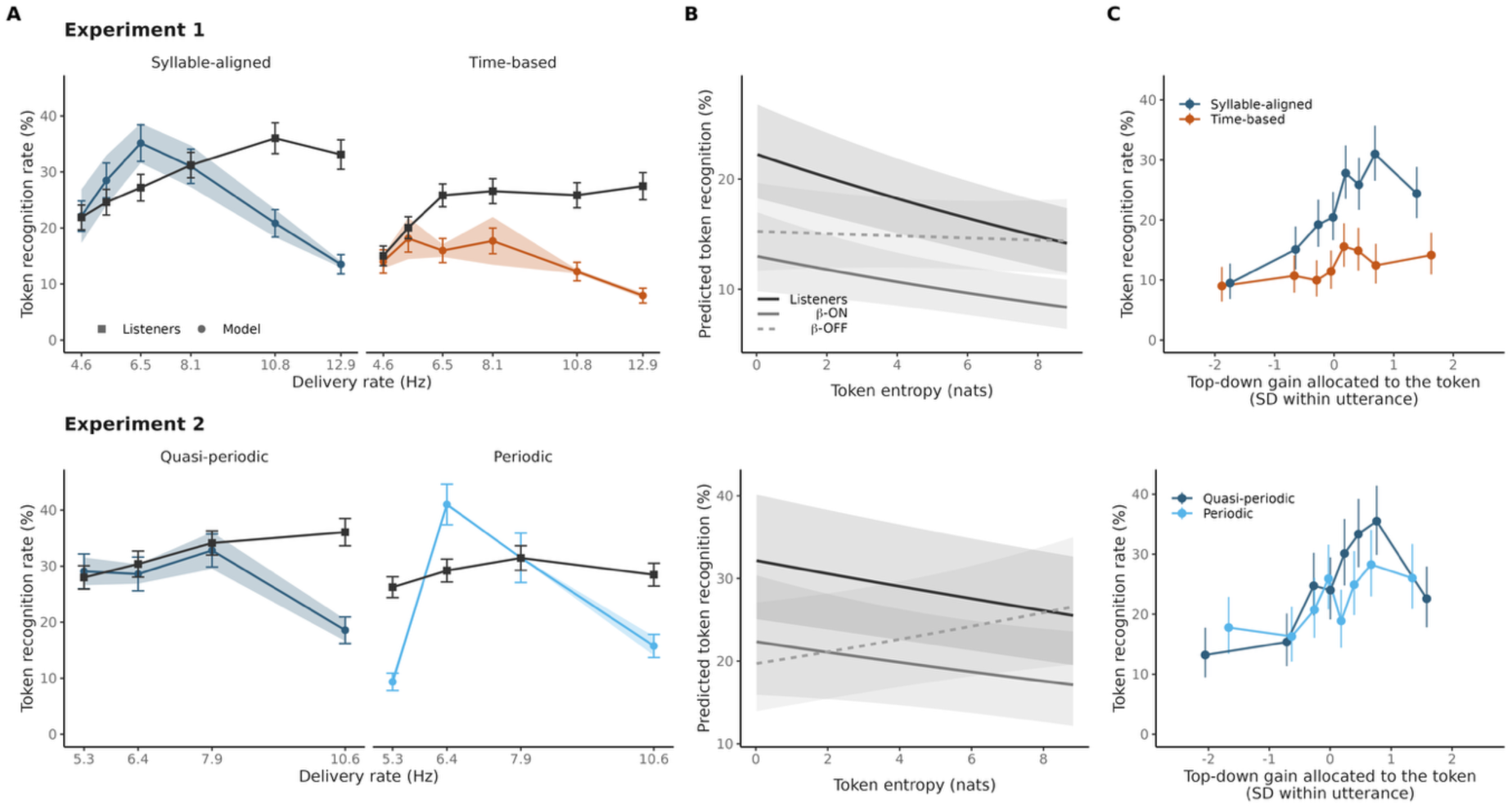
β-BRyBI compared with the listeners. Rows are the two experiments, columns the three analyses. (A) Token recognition rate against delivery rate for the listeners (black) and for the model (colors), by segmentation condition. Model points are the mean of β-ON and β-OFF (shown separately in Figure S3) and the band spans the two, so its width is the effect of beta modulation, not a confidence interval. Error bars are standard errors. (B) Token recognition predicted from token entropy for the listeners and for the two model states, with 95% bands, from one mixed model per experiment. Predictions are conditional on zero random effects, so levels are lower than the observed rates and the vertical scales differ between experiments; only slopes are comparable. (C) Token recognition rate against the top-down gain allocates to the token, within β-ON, in octiles of gain centered within the utterance. Error bars are binomial intervals. Colors follow Figures 2 and 4.

#### Only **β**-ON reproduces human sensitivity to contextual predictability

Listeners, *β*-ON and *β*-OFF are compared within a single fit per experiment, and the dependence of recognition on entropy is not the same across the three (*χ*^2^(2) = 7.78, p = .020 in Experiment 1; *χ*^2^(2) = 14.27, p < .001 in Experiment 2). In the listeners, a token is recovered less often the higher its entropy (slope = -0.062 log-odds per nat, 95% CI [-0.082, -0.042], z = 6.07, p< .001 in Experiment 1; slope = -0.037, 95% CI [-0.053, - 0.021], z = 4.52, p < .001 in Experiment 2). *β*-ON shows the same dependence (slope = -0.056, 95% CI [-0.095, -0.017], z = 2.80, p = .005 in Experiment 1; slope = -0.037, 95% CI [-0.080, 0.005], z = 1.72, p = .085 in Experiment 2), whereas *β*-OFF is flat in Experiment 1 (slope = -0.008, 95% CI [-0.044, 0.029], z = 0.42, p = .676) and runs the other way in Experiment 2, recognizing the least predictable tokens best (slope = 0.044, 95% CI [0.003, 0.085], z = 2.10, p = .035). *β*-OFF therefore departs from the listeners (difference = -0.055, 95% CI [-0.093, -0.016], z = -2.78, p = .016 in Experiment 1; difference = -0.081, 95% CI [-0.123, -0.039], z = 3.75, p < .001 in Experiment 2) while *β*-ON is indistinguishable from them (difference = -0.006, 95% CI [-0.048, 0.035], z = -0.31, p = .759 in Experiment 1; difference = 0.001, 95% CI [-0.043, 0.045], z = 0.03, p = .975 in Experiment 2, Figure 7B).

#### **β**-mediated prediction acts token by token, and its benefit depends on syllabic alignment

Within *β*-ON the top-down gain applied to a token varies from one token to the next, and tokens that received more of it are recovered more often (Experiment 1: *χ*^2^(1) = 63.90, p < .001; Experiment 2: *χ*^2^(1) = 34.70, p < .001). What the gain is worth depends on the units the segmentation delivers (Figure 7C). In Experiment 1 the relation is three times steeper where the boundaries fall on syllables (syllable-aligned: odds ration = 1.72 per standard deviation of gain, 95% CI [1.51, 1.96], z = 8.11, p< .001; time-based: odds ratio = 1.19, 95% CI [1.06, 1.34], z = 2.94, p = 0.003; interaction (*χ*^2^(1) = 17.23, p < .001). In Experiment 2, the regularity of the timing made no such difference (quasi-periodic: odds ratio = 1.34, 95% CI [1.18, 1.51], z = 4.71, p < .001; periodic: odds ratio = 1.40, 95% CI [1.18, 1.67], z = 3.90, p < .001; interaction *χ*^2^(1) = 0.22, p = .639). Permuting the gain across the tokens of a run removes the relation, so what predicts recognition is which token received the gain and not the overall amount of grain in that run.

## DISCUSSION

Speech unfolds as a continuous acoustic stream in which sensory evidence is inherently fleeting, yet comprehension remains stable across wide variations in speaking rate and acoustic clarity. Two classes of mechanism are usually invoked. The first is rhythmic alignment: theta-range dynamics (∼4–8 Hz) are thought to track the syllabic rhythm of speech and parse the stream into units amenable to linguistic analysis (Giraud & Poeppel, 2012; Gross et al., 2013; Peelle & Davis, 2012). The second is contextual prediction: internal models of linguistic structure generate top-down expectations about upcoming input and compensate when sensory evidence is degraded (Arnal & Giraud, 2012; Lewis & Bastiaansen, 2015; Ryskin & Nieuwland, 2023). The two accounts have developed in parallel, with limited empirical contact. By compressing speech threefold and then giving time back in controlled ways, the present experiment put them in contact and showed that they are not alternatives. Contextual prediction bears on recognition only where the temporal arrangement of the signal delivers syllable-sized units, and the computational model identifies beta-range modulation of the top-down pathway as what makes that dependence possible.

### Temporal compression as a probe of segmentation mechanisms

Compressing speech by a factor of three impairs intelligibility severely (Dupoux & Green, 1997; Pallier et al., 1998; Pefkou et al., 2017), and inserting silent intervals to lower the effective delivery rate partly restores it (Borges et al., 2018; Bosker & Ghitza, 2018; Ghitza & Greenberg, 2009). That restoration has been attributed to reinstating a theta-compatible delivery rate. The present results reproduce the compression and restoration effects and show that delivery rate alone does not explain them. At matched delivery rates, cutting at syllable onsets yields higher recognition than cutting every 62 ms, and the advantage holds at five of the six rates tested. What matters is therefore not only how fast the units arrive but also whether the boundaries coincide with syllabic units. This favors accounts in which theta dynamics flexibly align sampling to acoustic landmarks (Hovsepyan et al., 2020; Hyafil et al., 2015) over accounts that posit a fixed temporal scaffold.

### Recognition peaks above the canonical theta range

In both experiments recognition varied non-linearly with delivery rate and was best across a broad band from 8 to 13 Hz, above rather than within the canonical theta range. These are rates at which syllable-sized units arrive, not frequencies recorded from the brain, and they straddle the conventional boundary between the theta and the alpha band, so the optimum does not by itself say which neural band carries the parse. What it does speak against is a fixed temporal scaffold, since a sampling process that adapts to the material predicts an optimum that follows the signal, which a strict theta-tracking entrainment account does not. The segmentation advantage followed a related profile. It was smallest at 6.5 Hz, the only rate at which it did not reach significance, and largest at the two fastest rates. When speech arrives at a rate close to the natural syllabic one, it matters little where the cuts fall. The further the rate moves from it, the more their placement decides what is recognized. The canonical theta range is therefore better understood not as an optimal processing zone but as the range over which syllabic sampling proceeds unaided, with the best recognition falling just above it.

Listening outside that range is more demanding, and that cost probably leaves more than one trace. Alpha power rises with listening demand and has been proposed as an index of effort (Simeone et al., 2025), while alpha desynchronization over temporal cortex predicts correct recognition and has been read as a top-down enhancement of sensory processing (Dimitrijevic et al., 2017). Effort of that kind plausibly accompanies sampling at the edge of its range, and our data give no reason to set it against the account proposed here. What effort cannot explain is the effect of boundary placement: at a matched delivery rate it should be the same whether the cuts fall at syllable boundaries or at fixed intervals, yet recognition is not.

### Syllabic alignment is not sufficient: the spacing must follow the syllables

Experiment 2 holds alignment constant, every cut falling on a syllabic onset, and varies only the spacing of the intervals. If regularity were beneficial itself, isochrony should support recognition at least as well as pacing that follows the syllables (Kösem et al., 2018; Solli et al., 2025). The opposite is observed, and the cost of isochrony grows with delivery rate: quasi-periodic and periodic pacing do not differ at 5.3 and 6.4 Hz, differ at 7.9 Hz, and differ markedly at 10.6 Hz. Temporal regularity is therefore not a sufficient organizing principle for segmentation, and its cost appears exactly where the delivery rate leaves the range in which sampling proceeds unaided. A comparable dissociation has been reported outside the auditory domain, where predictions formed about the content of an upcoming event modulate perception independently of whether its timing is predictable (Lubinus et al., 2022). This is consistent with proposals that speech parsing operates through adaptively quasi-rhythmic rather than fixed oscillatory processes (Hyafil et al., 2015; Pittman-Polletta & Dilley, 2023). Taken with Experiment 1, effective segmentation requires two jointly necessary conditions: boundaries on syllables and intervals that follow those syllables.

### The usefulness of contextual prediction depends on the units segmentation delivers

Predictability bears on recognition only where segmentation returns syllable-sized units. In Experiment 1, the entropy effect is confined to syllable-aligned delivery, and in Experiment 2 to quasi-periodic pacing. This sets a boundary on what contextual prediction can do. Models of predictive language processing treat prediction as graded and simultaneously active at several levels of representation (Kuperberg & Jaeger, 2016; Heilbron et al., 2022). Much of the evidence for them concerns how strong and how specific predictions are (Pickering & Gambi, 2018; Dave et al., 2021; Ryskin & Nieuwland, 2023). Our results speak to a different aspect, the representation a prediction is applied to. A prediction can sharpen a representation only when one is available to be sharpened (Kok et al., 2013; Tuennerhoff & Noppeney, 2016), and in continuous speech we take that representation to be the syllabic parse. When delivery is imposed on a clock, the parse the listener obtains may not be one that lexical constraint can index, which would explain why the same predictability leaves recognition unchanged.

Experiment 2 provides the positive control for this dissociation: alignment is held constant across its two conditions, so the absence of an entropy effect under periodic pacing cannot be attributed to a failure of alignment. The entropy effect is present under quasi-periodic pacing and absent under periodic pacing. Natural speech is quasi-periodic rather than isochronous (Bosker & Ghitza, 2018; Ghitza & Greenberg, 2009), and strict isochrony removes the local timing variability that signals prosodic structure and phrase boundaries (Zhang et al., 2021; Pittman-Polletta & Dilley, 2023). Syllabic alignment on its own is therefore not sufficient. Contextual constraint appears to influence recognition only when the units’ segmentation delivers are the units over which that constraint is defined.

### **β**-band gating reproduces the listeners’ sensitivity to predictability

The computational modelling extends the behavioral findings by identifying a mechanism through which word-level prediction is timed relative to syllabic sampling. Enabling β-mediated lexical prediction in the β-BRyBI model does not make it a more accurate recognizer, but it changes the profile of sensitivity. With the modulation enabled, recognition follows contextual predictability as it does in the listeners, and with it disabled that relation disappears. This goes beyond prior demonstration in language processing (Arnal & Giraud, 2012; Bouton et al., 2018; Hovsepyan et al., 2023; Lewis et al., 2015; Zioga et al., 2023), by showing that a single mechanistic parameter, the gating of word-level predictions by a beta rhythm, is what carries a human-like dependence on context.

The reconfiguration is selective in a way that is theoretically informative. Within β-ON, the strength of the top-down prediction varies from one moment to the next, are recognition follows it. Indeed, units that arrive when it is high are recognized more often and this relation is more than three times stronger under syllabic alignment than under time-based segmentation. The loop releases its predictions in both cases, but they contribute to recognition far more when the boundaries fall on syllables. These findings converge with predictive-processing accounts in which the influence of a top-down prediction on sensory processing is weighted by its precision (Friston et al., 2020) and with the proposal that the feedback channel carrying that influence operates in the beta range (Arnal & Giraud, 2012; Bastos et al., 2015). They add a representational constraint. The beta rhythm sets when a prediction is released, while the parse sets whether there is anything for it to be released onto. Contextual influence requires both, which is why the same rhythm buys little when the units delivered are not syllables. In the model, the relation between top-down gain and recognition is three times steeper under syllabic alignment.

### Limitations and open questions

Several aspects of the present findings warrant further investigation. First, entropy at a given position is conditioned on everything that precedes it in the sentence, so the measure assumes a listener who carries the whole of that context. It describes an ideal observer, whereas the effective integration window of a listener is unknown and my itself vary with delivery rate. Second, the study draws on a constrained corpus of sentences from TIMIT, and it remains to be determined how the observed rate- and segmentation-dependent interactions generalize to more naturalistic speech, including prosodically variable and syntactically complex materials. Third, while the β-BRyBI model provides a principled computational account of the behavioral patterns, its predictions regarding the neural signatures of β-ON versus β-OFF, and particularly the expected contrast in beta-band power and its coupling to theta dynamics during syllabic versus time-based segmentation, have yet to be tested empirically. Such tests would provide a crucial link between the behavioral findings, the computational model, and the neural mechanisms proposed to implement beta-mediated lexical prediction.

Taken together, the present findings reframe the relationship between temporal structure and beta-mediated lexical prediction in speech comprehension. The two mechanisms are neither competing explanations for the same phenomenon nor simply additive contributors. Their contributions depend on one another, and time compression, by removing the temporal cues the parse ordinarily relies on, is what renders that dependence visible. Syllabic alignment is a structural prerequisite: it determines whether the word-level predictive system meets the input at the level over which it operates. Delivery rate determines whether that access carries a behavioral consequence: contextual predictability leaves no measurable signature in the middle of the canonical theta range, where syllabic sampling proceeds unaided and where boundary placement also matters least, and it acts on recognition at rates away from it. That range is therefore better understood not as an optimal processing zone but as the zone of spontaneous syllabic alignment, within which recognition proceeds without measurable help from context and outside which contextual predictability becomes decisive, though only when the parse has delivered syllables. Rather than passively synchronizing to the acoustic signal, the brain actively infers linguistic structure to stay ahead of incoming input, and the beta-band gating of those predictions is what tunes the tempo of speech segmentation.

## METHODS

### Ethic Statement

This study was approved by the Inserm Ethics Committee (Approval no. 20-689ter, August 31, 2023) and conducted in accordance with the Declaration of Helsinki. Informed consent was obtained from all participants.

### Participants

Fifty native speakers of American English completed Experiment 1 (26 female, 23 male, 1 undisclosed; mean age = 39.3 years, SD = 9.4) and sixty completed Experiment 2 (31 female, 29 male; mean age = 44.6 years, SD = 11.6). All reported normal hearing and no history of neurological disorders. Participants were recruited via Prolific (https://www.prolific.com/). Eligibility required being a native speaker of American English who used English on a daily basis, and completing the study on a laptop or desktop computer running Firefox. Full screening criteria are given in the Supplementary Materials. Participants took part in only one of the two experiments. Each received £6 for a session of approximately 40 minutes.

### Stimuli

Sentences were drawn from the TIMIT corpus (Garofolo et al., 1993). In both experiments, every experimental sentence was time-compressed by a factor of three using the WSOLA algorithm (Roelands & Verhelst, 1993); the calibration sentences were presented unaltered. Syllable boundaries were obtained with the automatic alignment method of Obin and Belião (2018). The median syllable duration was 187 ms in the original recordings, that is approximately 62 ms after compression, corresponding to a syllable rate of 16.1 Hz.

#### Experiment 1

From an initial pool of 125 sentences, 95 were retained so as to be matched for duration, syllable count, and prosodic balance. They ranged from 5 to 9 words (mean = 7.2, SD = 1.2) and from 6 to 16 syllables (mean = 10.4, SD = 2.0), comprised 2 to 6 prosodic phrases (mean = 3.7, SD = 1.0), and lasted between 1420 and 3756 ms (mean = 2558 ms, SD = 526 ms). Across the 95 experimental sentences, 86 speakers are represented, 28 female and 58 male, covering all eight dialect regions of the corpus, with nine speakers contributing more than one sentence (Table S1). Seven further sentences, drawn from outside this set, were presented uncompressed to every participant and provide the reference against which the effect of three-fold compression is measured (Figure 2A).

Two segmentation types were implemented. In the time-based condition, sentences were divided into fixed 62 ms segments. In the syllable-aligned condition, segments were defined by the syllable boundaries. In both conditions, a silent interval was inserted after each segment, scaled by a factor of 0.25, 0.5, 1, 1.5, 2, or 2.5 relative to the duration of the segment it followed, producing six delivery rates: 12.9, 10.8, 8.1, 6.5, 5.4, and 4.6 Hz. Since the segment is a fixed 62 ms under time-based segmentation and the syllable itself under syllable-aligned segmentation, these rates are matched on average across segmentation types rather than identical trial by trial.

Crossing the two segmentation types with the 6 delivery rates gives twelve conditions. A thirteenth condition presented the sentence compressed by a factor of three with no silence inserted, providing a baseline at the compression rate itself. Sentences rotated across these thirteen conditions, seven per condition.

#### Experiment 2

Ninety sentences were chosen for their proximity to the corpus mean in duration (M = 2730 ms, SD = 170 ms, range 2336 to 3078 ms), number of syllables (M = 11.3, SD = 0.8, range 10 to 13), and number of words (M = 8.1, SD = 0.7, range 7 to 9). They were spoken by 77 speakers, 26 female and 51 male, representing multiple dialect regions of the corpus DR1 4, DR2 10, DR3 19, DR4 13, DR5 11, DR6 10, DR7 19, DR8 4), with thirteen speakers contributing more than one sentence (Table S1). Four further sentences, drawn from outside this set, were presented uncompressed to every participant.

Segmentation was performed at syllable boundaries in every condition, each sound segment corresponding to one syllable. Two factors were manipulated, temporal regularity and delivery rate. In the quasi-periodic condition, the silence following each syllable was proportional to that syllable’s compressed duration, scaled by a factor of 0.5, 1, 1.5, or 2, so that natural inter-syllabic variability was preserved. The periodic condition was built to match it: the package duration was fixed at the median package duration of the corresponding quasi-periodic condition, that is 189, 157, 126, and 94 ms, so that syllable onsets occur<u>r</u>ed at uniform intervals. The two regularity conditions therefore differ in the variability of the intervals between syllable onsets while sharing the same median package duration, and <u>are</u> both described by the same nominal delivery rates of 5.3, 6.4, 7.9, and 10.6 Hz.

### Procedure

The experiment was programmed in PsychoPy and deployed online via Pavlovia (https://pavlovia.org/). On starting, participants were randomly assigned to one of the stimulus lists. In Experiment 1 each participant completed 102 trials: 95 compressed sentences distributed over 13 blocks, the twelve experimental conditions and the fully compressed baseline, and seven uncompressed sentences. In Experiment 2 the set comprised 90 compressed sentences over nine blocks and four uncompressed sentences. Block order and trial order within blocks were pseudorandomized. Short breaks were permitted between blocks.

Participants were instructed to complete the study in a quiet environment using stereo headphones. Headphone use was established by self-report at recruitment, participants being eligible only if they confirmed having a computer and headphones available for the study, and by explicit instruction at the start of the session. Participants were asked to adjust the volume to a comfortable level and were reminded to avoid external distractions. The task could only be accessed from a desktop or laptop computer running the Firefox browser to ensure compatibility and stable audio playback.

The session began with the uncompressed sentences, seven in Experiment 1 and four in Experiment 2. They served to verify engagement and exclude non-compliant participants and they provide the uncompressed reference against which the effect of compression is measured; they therefore occupy the opening of every session. This was followed by five practice trials, each with a 15-second response window. Practice trials and uncompressed sentences were drawn from outside the experimental set. Each trial proceeded as follows: (i) a 1000 ms fixation cross, (ii) binaural presentation of a stimulus sentence, (iii) a 15-second response for typed recall, (iv) a completion screen, and (v) a 1000 ms inter-trial interval. Stimulus presentation and response screens were time-locked and non-skippable. The session lasted approximately 40 minutes, depending on individual response times and break duration.

### Scoring

Responses were scored at the token level, the unit over which GPT-2 assigns probabilities and therefore the unit in which contextual entropy is defined. A token is an entry in the sub-word vocabulary produced by byte-pair encoding, which merges the character sequences recurring most often in a training corpus into single entries. The merging is orthographic and frequency-based rather than phonological: frequent words form a single token whatever their length, “always” and “before” as much as “the”, while rarer words are split into two or three. Tokens are accordingly intermediate in grain between words and syllables, and closer to words: the 95 experimental sentences contain 679 words, 771 tokens and 971 syllables. Scoring at this level puts the recognition measure and the contextual entropy measure in the same units. The tokenization depends only on the text, so a sentence yields the same tokens in every condition and for every participant.

### Scoring was fully automatic

The typed response and the reference sentence were lowercased, stripped of punctuation, and encoded into tokens; the two token sequences were aligned by longest common subsequence, and a reference token was counted as recognized only when the alignment matched it to an identical token in the response. The trial score is the proportion of reference tokens recognized. The criterion is orthographic: typographical errors, alternative and compound spellings, and homophones of the target all count as errors, as do substitutions that fit the preceding context, since the target is the word the talker produced and contextual plausibility plays no part in the criterion. Trials on which the participant typed nothing were scored as zero tokens recognized; there are 253 such trials out of 5100 in Experiment 1, and none in Experiment 2.

### Sample Size Calculation

The target sample was set before data collection on the basis of an effect size of R² = .30 reported by Ramus et al. (2021, unpublished data) in a comparable regression model. Converting this to Cohen’s yielded f² = 0.43, an a priori power analysis using G*Power (Faul et al., 2009) for a linear model with three predictors, *α* = 0.05, and desired power of .95 returned N = 45. Increasing this by approximately 10% to allow for exclusions and data loss gave a target of 50, the number recruited for Experiment 1. This calculation fixed the number of participants to recruit.

### Statistical analysis

All models were generalised linear models fitted with glmmTMB (version 1.9.17, Brooks et al., 2017) in R (version 4.5.2); marginal means and contrasts were obtained with emmeans (version 2.0.0), and omnibus tests with car (version 3.1-3).

Two levels of analysis are used. The effects of compression, of segmentation, and of delivery rate take the trial as the unit: the response is the proportion of the sentence’s tokens correctly reported on that trial, fitted with a beta family and a logit link, with random intercepts for participant and sentence. A beta response is defined on the open unit interval, so proportions of exactly zero or one were shifted inward by 10^−4^. The effects of contextual entropy take the token as the unit: each token is scored as recovered or not, fitted with a binomial family and a logit link, with random intercepts for participant, sentence, trial and token position. Token positions 2 to 11 enter the entropy analyses, the first token of a sentence having no preceding context over which entropy is defined, and positions beyond the eleventh occurring in too few sentences to support a cell.

Each part of the Results rests on a single model. Delivery rate is a categorical factor with treatment coding and the slowest rate as reference. Type III omnibus tests of the main effects are computed under sum-to-zero contrasts, since under treatment coding a Type III main effect is evaluated at the reference level of the other factor rather than averaged over it; the contrasts reported are estimated from marginal means and do not depend on the coding.

Every test of a term within a fitted model is a Wald chi-square test. Every comparison between two nested models is a likelihood ratio test. The latter are used three times in Experiment 1 and four times in Experiment 2: for the effect of three-fold compression, for the quadratic term of the delivery-rate profile, for the interaction between the shape of that profile and the manipulated condition, and, in Experiment 2 only, for the quadratic term restricted to the manipulated rates. Families of contrasts estimated one per delivery rate, or one per cell, are corrected with the Benjamini-Hochberg procedure; the pairwise comparisons among the three conditions of the baseline model use Tukey’s adjustment.

#### Experiment 1

All 5100 trials are analysed. The effect of three-fold compression is a likelihood ratio test between the model with and without the condition term, and the difference is reported on the response scale. The delivery-rate profile is fitted as a quadratic over the seven rates from 4.6 to 16.1 per second, the fully compressed condition being the fastest of them; the quadratic term is tested against the linear model by likelihood ratio, as is the interaction between segmentation type and the shape of the profile. Segmentation type and delivery rate then enter one Segmentation × Delivery rate model, from which the main effect of segmentation averaged over rates, the main effect of rate, their interaction, and the six per-rate contrasts are all read. The fully compressed condition carries no delivery rate and cannot enter that design, so each segmentation type is compared with it in a separate three-level model. Contextual entropy enters an Entropy × Segmentation × Delivery rate model over 29 614 tokens, the twelve per-cell slopes being corrected together.

#### Experiment 2

5617 trials are analysed. The structure is that of Experiment 1, with temporal regularity in place of segmentation type and four delivery rates from 5.3 to 10.6 per second, the fully compressed condition again entering the rate profile at 16.1. The quadratic term is tested twice, over the full profile and over the manipulated rates alone. Regularity and delivery rate enter one Regularity × Delivery rate model; the fully compressed condition is compared with each regularity regime in a separate three-level model; and contextual entropy enters an Entropy × Regularity × Delivery rate model over 39 519 tokens, the eight per-cell slopes being corrected together.

Entropy Scores. Contextual uncertainty was quantified with the GPT-2 language model, accessed via the Hugging Face Transformers library. Byte-pair encoding gives GPT-2 a fixed inventory of 50,257 sub-word entries, its vocabulary, and the model represents any text as a sequence of entries drawn from it. At each position the model returns a probability for every entry in that inventory, the 50,257 values summing to one. Entropy was computed as the Shannon entropy of that distribution, in nats:

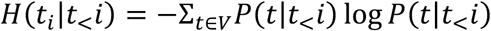

where *V* is the model’s vocabulary and *P*( *w* ∣ *w*_<*i*_) denotes the probability assigned to token *t* given the context of preceding tokens. One value is obtained per token, and no aggregation to the word was performed: the token is the unit at which the model defines probability and the unit at which responses are scored, so entropy and recognition are measured on the same units throughout.

The first token of a sentence has no preceding context and carries no entropy, so analyses begin at the second. Entropy depends only on the sentence, not on how it was presented, so a given token carries the same value in every condition and for every participant. Across the stimulus set, the values range from 0.00 to 8.83 nats, against a ceiling of log (50,257) = 10.82 nats for a uniform distribution over the vocabulary.

Entropy is a property of the context, not of what any listener heard. It is therefore computed from the intact sentence, which assigns the same value to every participant at a given token position and keeps the predictor independent of the outcome being modelled. A listener who misses part of a sentence has a poorer context at the next token, and the entropy that listener experiences is higher than the value used here. That difference depends on how much of the sentence was recognized and not on delivery rate (Figure S2), so it is not aligned with the contrasts the analysis tests.

### Computational Model

Rhythms are generated by the Wilson-Cowan model with SNIC bifurcation:

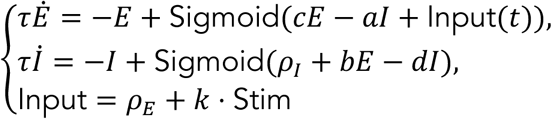

where *E*(*t*), *I*(*t*) are excitatory and inhibitory population activity, *k* is the coupling strength to the normalized stimulus envelope Stim(*t*), and *ρ_E_* is a background input.

The *γ*-rhythm is defined via CFC to the phase of the theta rhythm *τ_θ_*:

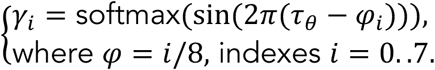

Words are represented as matrices of syllables: 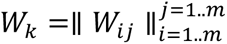, where *m* is the number of syllables in the dictionary. *W_i_*_j_ = 1 if *j*-syllable follows *i*-syllable, else *W_i_*_j_ = 0. The word-matrix is defined as the weighted sum of all words and phrases: 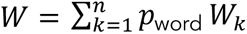, *n* is a number of words in the dictionary.

The linguistic part of *β*-BRyBI can be described as an interactive activation model for syllables that is temporally controlled by theta rhythm. Decoded from the spectrum, phonemes activate the corresponding elements in the network of syllables. Concurrently, word-level generates a pattern of possible transitions between syllabic elements:

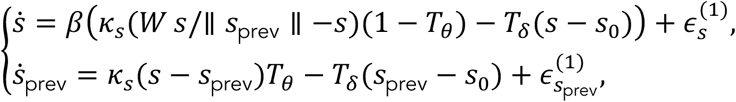

where *s* and *s*_prev_ are two -syllable pointers, and *T*_ẟ_ is a trigger that switches words. Beta rhythm (*β_freq_* = 17 *Hz*) modulates the usage of information from the top level for the inference of syllables in betaON regime and *β* = 1 in the case of betaOFF regime.

The auditory spectrogram is formed via the convolution of syllables and phonemes with predefined tensors for each frequency band *P_fγθ_*:

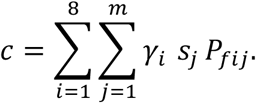

Inference of this generative model was done by using Dynamical Expectation-Maximization algorithm with SPM12 matlab library.

### Statistical Analysis

Only model runs on the recordings the listeners actually heard enter the analyses, and the fully compressed condition, which has no delivery rate, is excluded throughout. Model output and listeners’ responses are read at the same level, that of the byte-pair token, and the first token of each sentence, which has no preceding context, is excluded on both sides. All models are binomial generalized linear mixed models fitted with glmmTMB, with crossed random intercepts for sentence, for token position and for run, a run being one presentation of one sentence to one listener or one pass of the model over one sentence. Delivery rate enters as a categorical factor. Omnibus tests are Type III Wald chi-square tests computed under sum-to-zero contrasts, and contrasts estimated from marginal means are corrected within their family with the Benjamini-Hochberg false-discovery-rate procedure.

Four analyses are reported, each a single model per experiment.

The first asks whether the experimental manipulations act on the model as they act on the listeners. Token recognition in the model is modelled on segmentation condition, delivery rate and model state (β-ON, β-OFF), with all their interactions.

The second asks how far the model departs from the listeners, and where. Token recognition is modelled on Source (listeners or model), segmentation condition, and delivery rate, with all their interactions. The difference between the model and the listeners is estimated in each segmentation by delivery rate cell, and these differences are then combined by planned linear contrasts on the same fit into a theta band, at or below 8 Hz, and a band above it, so that the comparison between bands is a contrast and not a separate model fitted on a subset.

The third asks whether the two states of the model reproduce the listeners’ sensitivity to contextual predictability. The three sources, listeners, β-ON and β-OFF, enter as a single three-level factor, and token recognition is modelled on token entropy in interaction with that factor, with segmentation condition and delivery rate as additive terms. Entropy is in nats and centred on its grand mean, so that a slope is the change in log-odds of recognition per nat. The three slopes and the three differences between them are contrasts on this one fit.

The fourth asks what the beta gating does within the state that has it. In β-ON, each token carries the mean top-down gain applied over its window. This gain is centred within <u>the</u> run, so that every comparison is between tokens of the same utterance in the same condition, and scaled by its within-utterance standard deviation, so that the two experiments share a unit. The analysis covers token positions 2 to 11, and three tokens out of some fourteen thousand whose gain exceeds the theoretical maximum of one are excluded as numerical divergences of the loop. Token recognition is modelled on the centered gain in interaction with segmentation condition and delivery rate.

## Supporting information

Supplementary Materials for the Manuscript

## ACKNOWLEDGEMENTS

We gratefully acknowledge support by the Fyssen Foundation (PI: S.B.), by the Fondation pour l’Audition (RD-2016-R to S.B. and FPA IDA11 to A.L.G), by Agence Nationale de la Recherche (ANR-21-CE28-0028 to S.B.), by Evolving Language, Swiss National Science Foundation Agreement #51NF40_180888 (to A-L.G.). This work has also benefited from a French government grant managed by the Agence Nationale de la Recherche under the France 2030 program, reference ANR-23-IAHU-0003. We are grateful to Boris Gutkin for making this work possible through their institutional support.

## AUTHOR CONTRIBUTIONS

S.B. and A.L.G. led the conceptualization of the project and coordinated the research program. O.P. and S.B. designed the experiments. O.P. implemented the experimental paradigm, conducted data collection, and managed data curation. O.P., O.D., and S.B. developed the experimental and analytical software. O.P. and S.B. performed the statistical analyses. A.L.G. and O.D. contributed to the development of the computational modeling framework. O.D. developed the computational model and conducted the associated simulations. O.P. and O.D. drafted the original manuscript and prepared the figures. S.B. reviewed and edited the manuscript and contributed to figure refinement. A.L.G. provided theoretical guidance and contributed to manuscript revision. S.B. and A.L.G. supervised the project and secured funding. All authors approved the final version of the manuscript.

## DATA AVAILABILITY STATEMENT

All data and analysis files are available from the Institut Pasteur GitLab repository at https://gitlab.pasteur.fr/sobouton/repack/-/tree/55c078e7d3a965cc11c8efa32df3255c7fa4bbe7/https://gitlab.pasteur.fr/ida-public/repack.

## CODE AVAILABILITY STATEMENT

Code for the beta-BRyBI model, analysis code for both experiments, and associated data are available at Zenodo https://doi.org/10.5281/zenodo.22079873 and from the Institut Paster GitLab repository at: https://gitlab.pasteur.fr/ida-public/repack.

