## Supplementary Materials for the Manuscript for "Contextual Prediction Tunes the Tempo of Speech Segmentation"

‡co-last authors

### Stimuli

| | Experiment 1 | | | Experiment 2 | | |
|---|---|---|---|---|---|---|
| Dialect region | Male speakers | Female speakers | Total number | Male speakers | Female speakers | Total number |
| DR1: New England | 3 | 1 | 4 | 3 | 0 | 3 |
| DR2: Northern | 10 | 5 | 15 | 7 | 2 | 9 |
| DR3: North Midland | 11 | 8 | 19 | 10 | 6 | 16 |
| DR4: South Midland | 16 | 1 | 17 | 10 | 2 | 12 |
| DR5: Southern | 3 | 8 | 11 | 2 | 7 | 9 |
| DR6: New York City | 5 | 2 | 7 | 4 | 4 | 8 |
| DR7: Western | 9 | 1 | 10 | 12 | 4 | 16 |
| DR8: Army Brat | 1 | 2 | 3 | 3 | 1 | 4 |
| Total | 58 | 28 | 86 | 51 | 26 | 77 |

***Table S1*. Distribution of TIMIT speakers by dialect region and sex.** Entries are numbers of unique speakers contributing experimental stimuli.

### Power Analysis

The target sample size was set before data collection, from statistical parameters reported by Ramus et al. (2021, un published data), who fitted multiple regression models with quadratic terms to speech processing data and reported a coefficient of determination of $R^2 = 0.30$ for the relationship between interval length and the outcome of interest. Converting that value gives Cohen's $f^2$:

$f^2 = \frac{R^2}{1-R^2} = \frac{0.30}{1-0.30} = 0.4286$, a large effect by conventional benchmarks.

We then computed the sample size required to achieve 95% power ($1 - \beta = 0.95$, at a significant threshold of $\alpha = 0.05$, for a model with three predictors including the linear and quadratic terms using the opion **"F tests - Linear multiple regression: Fixed model, R² deviation from zero"** in G*Power (Faul et al., 2009).

| Parameter | Value |
|---|---|
| **Effect size *f*²** | 0.4286 |
| **Significance level** | 0.05 |
| **Power** | 0.95 |
| **Predictors** | 3 |
| **Total sample size (computed)** | **45 articipants** |

The output parameters were a noncentrality parameter (λ) = 19.29, a Critical F-value of 2.83 on 3 and 41 degrees of freedom, and an actual power of 0.954. Increasing this figure by approximately 10 % to allow for exclusions and data loss gave a target of 50 participants. This calculation fixed the number of participants to recruit.

### Data collection

#### Prolific experiments.

The online experiment comprised two parts: the prescreening and experimental session. A brief questionnaire "Eligibility for Participation in a Speech Perception Study" was intended for Prolific participants who met specific inclusion criteria. Please note that we do not ask for this information directly; instead, we rely on the following Prolific's built-in participant filters:

- **First Language**: English
- **Nationality**: United Kingdom
- **Cultural Identity**: I identify as a *monocultural* individual
- **Primary Residence Before Age 18**: United Kingdom
- **Language-related Disorders**: None
- **Primary Language**: English
- **First Language Acquired in Life**: English
- **Dyslexia**: No
- **Neurodiversity**: No
- **Hearing Difficulties**: No
- **Vision**: Yes

Survey Details

**Title**: *Prescreening for a 40-minute experiment – BRyBI*

**Estimated time**: 1–2 minutes

This is a short prescreening survey for an upcoming 40-minute online experiment on speech perception. In the main study, you will listen to degraded audio recordings of sentences and type as many words or phrases as you can recognize.

If you meet the criteria and your responses align with our inclusion goals, you may receive an invitation to participate in the full experiment.

Thank you for your time and interest!

Prescreening Questionnaire consists of 4 questions:

1. Is English your first language? Yes/No
2. If you speak several languages, how much time do you spend speaking English in your daily life?
    - all the time: I do not speak other languages
    - all the time, 100%
    - 90% of the time
    - 80% of the time

- 60-70% of the time
- half of the time
- less than 40% of the time
- I barely use English at the moment

3. Do you have a computer and headphones to use for the online study? Yes/No
4. Do you have Mozilla Firefox browser? Yes/No

Participants who reported English to be their first language, that they use all the time, as well as reported to have a computer, headphones and Mozilla Firefox browser were invited to take part in the experimental session. Please note that we did not do additional headphone check but explicitly asked participants to use headphones: “IMPORTANT: Please use headphones and the Mozilla Firefox browser to assure the correct completion of the task and please make sure there are no external distractions for the next 40 minutes”.

**Supplementary Figures**

Figure S1

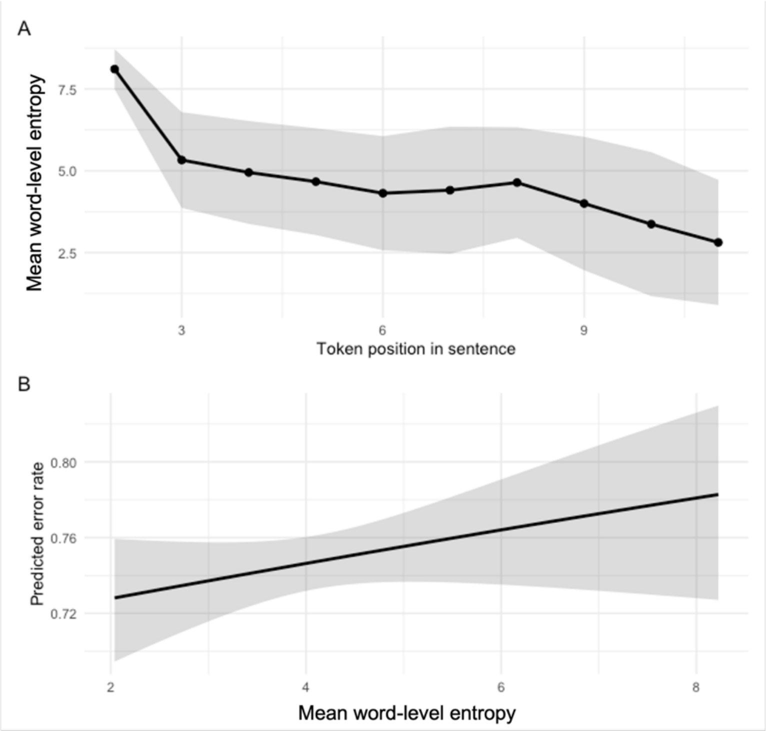

**Word-level entropy across sentence position and its effect on recognition accuracy. (A)** Mean word-level entropy (computed via GPT-2) as a function of token position within the sentence. Entropy is highest at sentence onset, reflecting maximal lexical uncertainty in the absence of prior context, and decreases progressively as contextual information accumulates across the sentence. Points represent mean entropy per token position; the shaded band indicates 95% confidence intervals. **(B)** Model-predicted error rate as a function of mean word-level entropy, collapsed across all segmentation conditions and delivery rates. Higher contextual uncertainty (higher entropy) is associated with increased predicted error rates, indicating that lexical predictability exerts a measurable influence on recognition under extreme temporal compression. The solid line shows the fitted beta mixed-effects model; the shaded band indicates 95% confidence intervals.

Figure S2

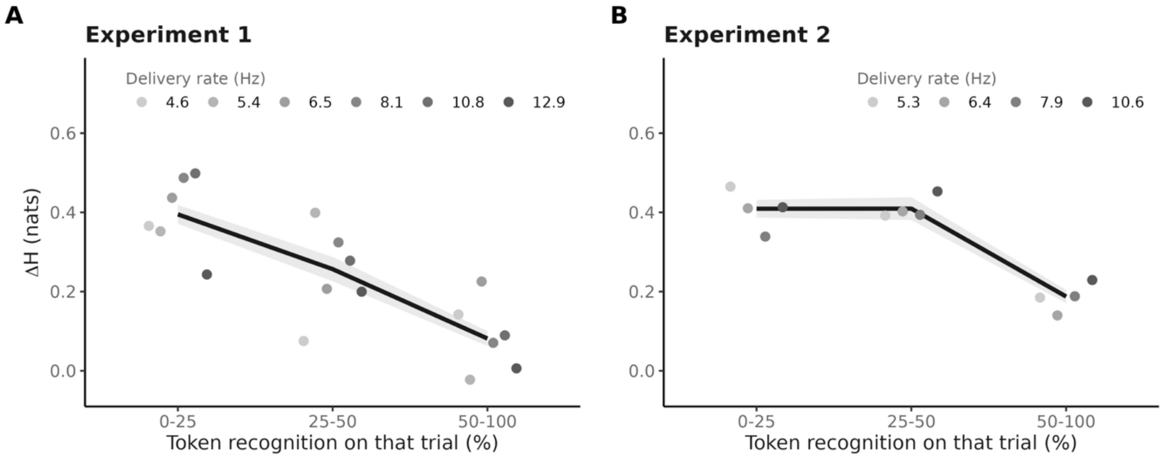

**Figure S2.** What determines the divergence between the listener's context and the intended one. For every trial, entropy was recomputed from the participant's own transcription, H(perceived), and compared with the value obtained from the intact sentence, H(true); $\Delta H = H(\text{perceived}) - H(\text{true})$. The black line is the relation between ΔH and the token recognition rate achieved on the same trial, pooled over delivery rates, with its standard error; the grey points are the same quantity computed separately at each delivery rate. (A) Experiment 1, six delivery rates. (B) Experiment 2, four delivery rates. ΔH falls as recovery improves, and the delivery rates scatter around that relation without ordering themselves: once recovery is accounted for, delivery rate contributes nothing to ΔH (Experiment 1: $\chi^2(5) = 3.72$, $p = .59$; Experiment 2: $\chi^2(3) = 2.34$, $p = .51$).

Figure S3

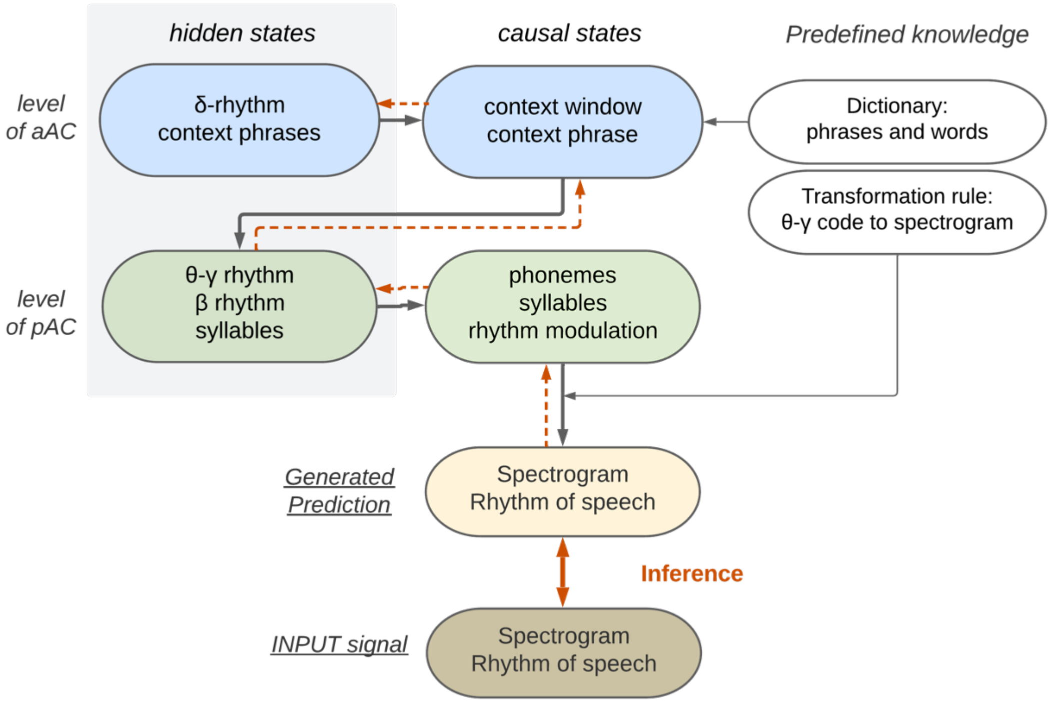

**Figure S3. Architecture of the β-BRyBI generative model.** The model operates as a hierarchical generative architecture organized across two cortical levels. At the level of primary auditory cortex (pAC; green), θ-γ and β rhythms drive the inference of phonemes and syllables from the incoming acoustic signal, with rhythm modulation coordinating the temporal dynamics of evidence accumulation. At the level of associative auditory cortex (aAC; blue), δ-rhythm dynamics maintain context phrases within a sliding context window, integrating phrase- and word-level representations over longer timescales. Predefined knowledge — comprising a dictionary of words and phrases and a transformation rule mapping θ-γ codes to spectrograms — constrains inference at both levels. Solid grey arrows indicate bottom-up (feedforward) message passing; dashed orange arrows indicate top-down (feedback) predictive signals. The model performs inference by minimizing the discrepancy between the internally generated spectral prediction (Generated Prediction) and the externally provided acoustic input (INPUT signal), as indicated by the bidirectional orange arrow labeled *Inference*. β-mediated prediction (β-ON) selectively modulates the propagation of top-down lexical expectations onto syllabic representations at the pAC level, enabling contextual prediction to compensate for degraded bottom-up temporal cues when syllabic segmentation is preserved.

Figure S4

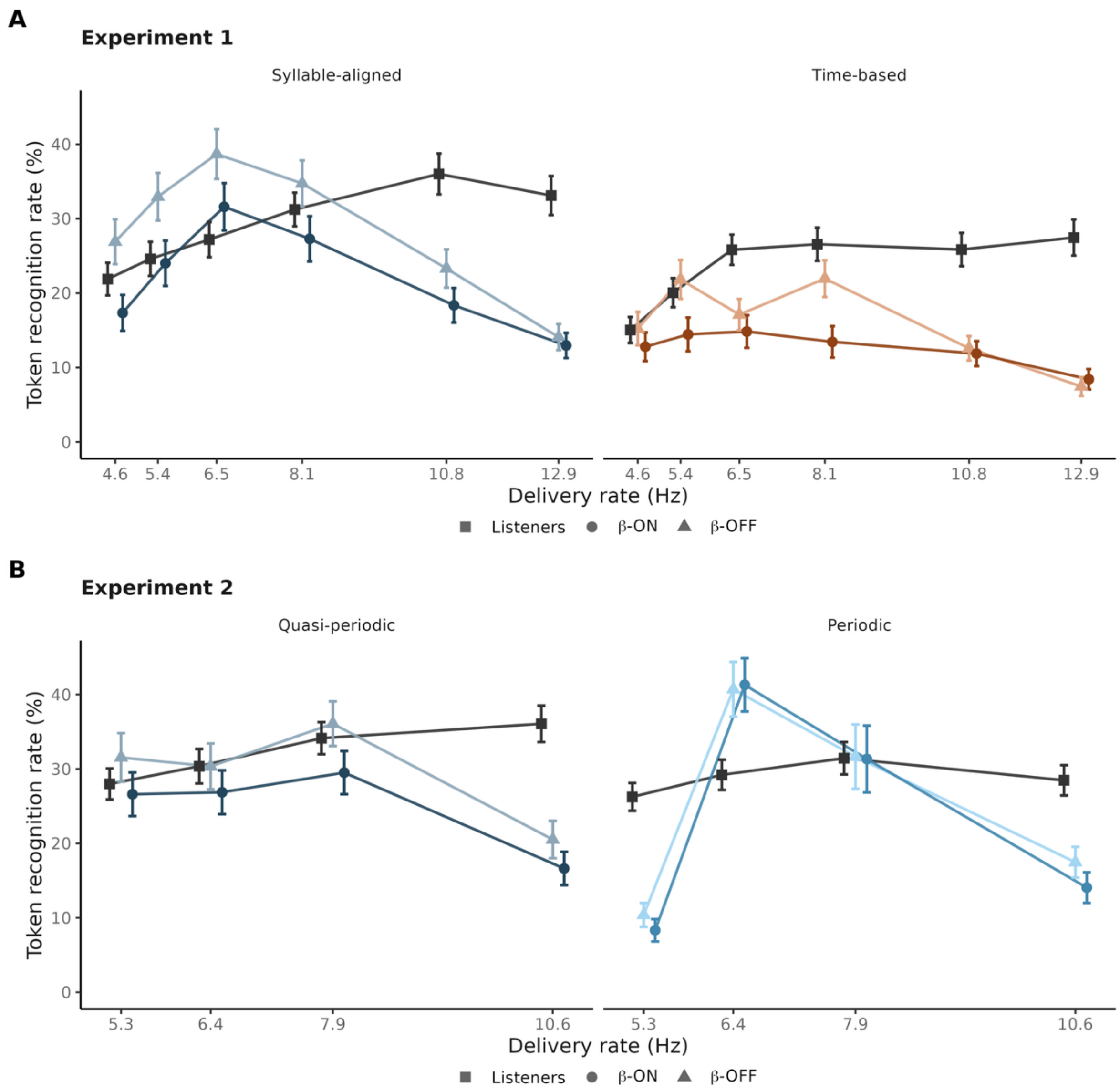

**Figure S4. β-ON and β-OFF shown separately.** Token recognition rate as a function of delivery rate, for the listeners (black squares), β-ON (circles) and β-OFF (triangles), by segmentation condition, in Experiment 1 (**A**) and Experiment 2 (**B**). β-ON carries the colour of its condition and β-OFF the same hue lightened. Error bars are standard errors over sentences for the two model states and over participants for the listeners. These are the same model data as in Figure 7A, where the two states are collapsed into a single series.
